# Genome-wide circadian gating of a cold temperature response in bread wheat

**DOI:** 10.1101/2022.11.29.518321

**Authors:** Calum A. Graham, Pirita Paajanen, Keith J. Edwards, Antony N. Dodd

## Abstract

Circadian rhythms coordinate the responses of organisms to their daily fluctuating environments, by establishing a temporal program of gene expression. This schedules aspects of metabolism, physiology, development and behaviour according to the time of day. Circadian regulation in plants is extremely pervasive, and is important because it underpins both productivity and seasonal reproduction. Circadian regulation extends to the control of environmental responses through a regulatory process known as circadian gating. Circadian gating is the process whereby the circadian clock regulates the response to an environmental cue, such that the magnitude of response to an identical cue varies according to the time of day of the cue. Here, we show that there is genome-wide circadian gating of responses to cold temperatures in plants. By using bread wheat as an experimental model, we establish that circadian gating is crucial to the programs of gene expression that underlie the environmental responses of a crop of major socioeconomic importance. Furthermore, we identify that circadian gating of cold temperature responses are distributed unevenly across the three wheat subgenomes, which might reflect the geographical origins of the ancestors of modern wheat.

**One-sentence summary:** There is genome-wide circadian gating of a response to low temperatures in a crop of major socioeconomic importance.

## Introduction

Circadian rhythms are biological cycles with a period of about 24 h that persist in the absence of environmental cues. A feature of circadian rhythms is that they determine the time of day (phase) when specific processes occur [1, 2]. The process of entrainment aligns the phase of the circadian clock with the phase of the environment and, in turn, the circadian clock extensively influences the phase of gene expression, metabolism and physiology [3].

In plants, one way that this occurs is through clock controlled gene promoters, which temporally coordinate transcript accumulation according to the time of day [1, 4–8]. The regulation of cellular processes by circadian clocks is important for crop performance, and understanding circadian regulation in crops forms an important part of the strategic design of climate change-resilient crops [9–12].

Fluctuating spring temperatures and unpredictable cold periods caused by climate instability represent a threat to cereal crops worldwide [13–15]. During multiple stages of development, wheat is susceptible to low temperature conditions [16–18], including during seedling establishment [19, 20]. Low temperatures trigger genome wide responses in wheat that can reduce photosynthetic capacity, cause photoinhibition, and consequently reduce yield [18, 21–25]. Therefore, wheat represents an excellent model to understand the processes that underlie responses of plants to low temperature conditions, because this has a direct impact upon crop yield.

One way that the circadian oscillator influences responses to environmental cues, such as temperature fluctuations, is through circadian gating. Circadian gating is the process whereby the circadian clock regulates a response to an acute environmental stimulus, such that the response occurs only at certain times of day, or the magnitude of the response varies according to the time of day that the stimulus occurs [26, 27]. In plants, there are a number of examples of the circadian gating of specific responses to high and low temperature cues [9, 28–36], with gating-like processes also occurring in cultivated rice fields and wild plant populations [37, 38]. Changes in the composition of the circadian-regulated transcriptome occurs under drought and low temperature conditions [32, 39–42], and there is circadian gating of heat stress responses [32, 40, 41]. However, we do not know whether the circadian gating of cold temperature responses occurs at a genome-wide scale in plants. This remains an important open question, because the pervasive nature of circadian gating of temperature responses makes it a core part of understanding how plants- including crops- respond to their fluctuating environments.

In hexaploid bread wheat, around 30% of the transcriptome is regulated by the circadian clock [43]. Circadian clock gene loci such as *TaPpd* (orthologous to Arabidopsis *PRR3* or *PRR7*) determine agriculturally-important photoperiod responses of domesticated wheat [44–47]. The complexity of this signal integration is increased in polyploid crops such as wheat by the presence of multiple gene paralogs or homoeologs that can have differential patterns of circadian regulation [39, 43]. In hexaploid wheat, over 50% of genes (approximately 50,000 genes) occur as sets of three gene homoeologs termed triads [48, 49], providing an interesting genomic structure to study environmentally-responsive gene expression within polyploid genomes.

We investigated the involvement of circadian gating in the genome-wide responses of hexaploid bread wheat to low temperature. Given that the cold-responsive CBF pathway is gated by the circadian clock in Arabidopsis [28] and a third of wheat genes are circadian- regulated [43], we reasoned that there might be a pervasive influence of the circadian clock upon transcriptional responses to short periods of low temperature. Using wheat as a model to investigate this question provides insights into a species of vast socioeconomic importance, and more generally the integration of circadian and environmental cues in polyploids. Taken together, our data identify a major role for circadian regulation in shaping the cold-responsive transcriptome in plants.

## Results

### The time of day affects the number of genes that respond to cold temperature conditions

We wished to discover whether there is circadian gating of responses to acute cold temperature cues in hexaploid bread wheat (*Triticum aestivum)*. We used low temperature treatments as an experimental model because circadian gating of responses to short cold treatments has been reported for specific Arabidopsis genes [28, 50, 51]. It is not known whether there is circadian gating of low temperature responses across the entire genome in plants. Spring wheat varieties are more cold-sensitive than winter wheat [52, 53], and are exposed to low spring temperatures during seedling establishment. Therefore, for our study, we selected a cold-responsive hexaploid spring wheat cultivar (Cadenza) that has no vernalization requirement, some low temperature tolerance, and good genome sequence coverage and germplasm resources.

To test for the occurrence of circadian gating, short cold treatments were applied to separate batches of wheat seedlings under free running (constant) conditions (Fig. 1A). This comprised identical three-hour cold treatments (4°C), at six sequential timepoints, following transfer to free running conditions (Fig. 1A). This design allows the detection of transcripts that have a response to cold that is restricted to certain times of day, or a 24 h fluctuation in sensitivity to the cold treatment. After each treatment, tissue was harvested for RNA isolation from the second leaf of control temperature and cold-treated seedlings, with each seedling sampled once only (Fig. 1A). Transcript count values (isoforms) were condensed to the gene level, so transcript abundance changes described here refer to changes at the gene level.

**Figure 1.**
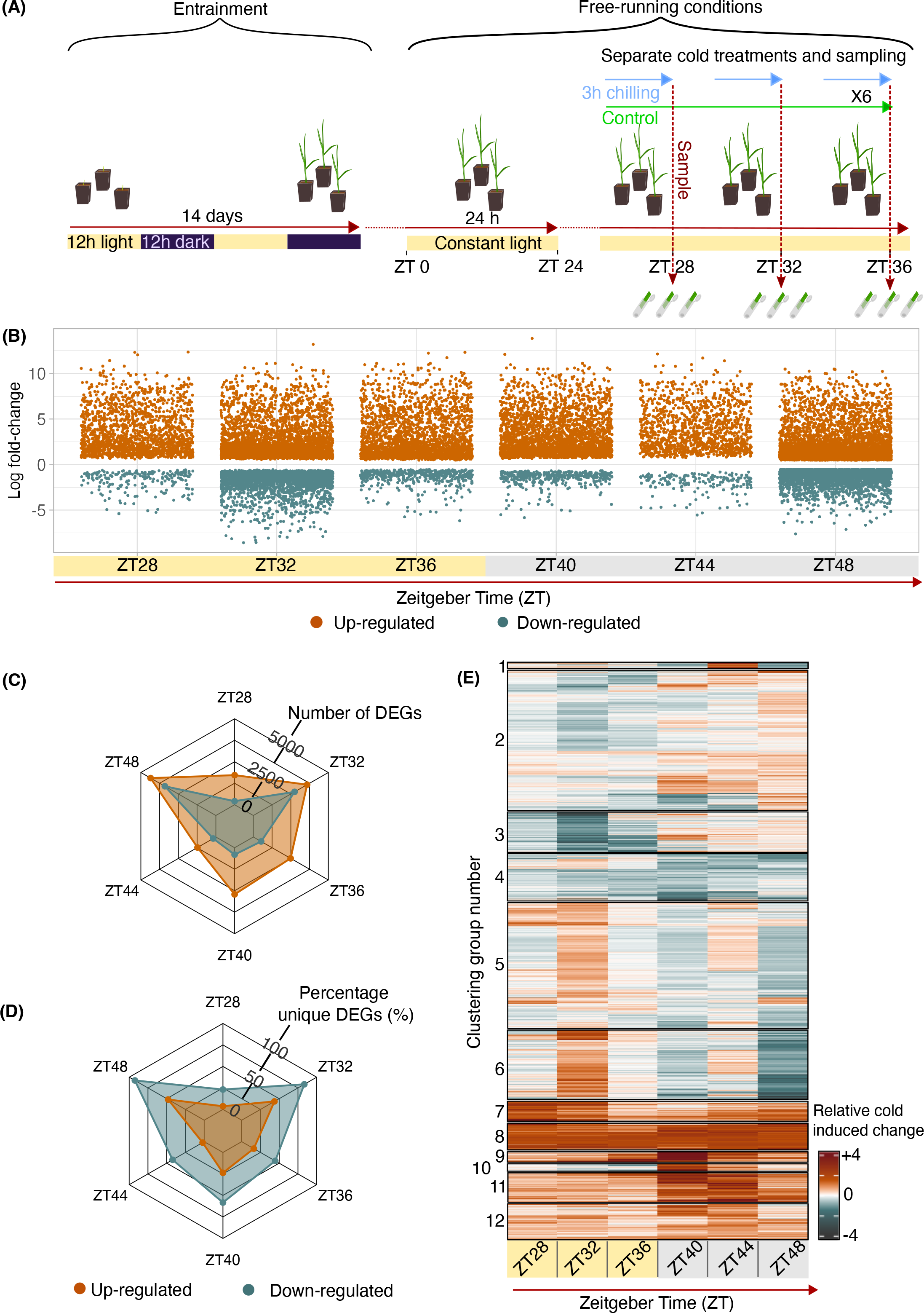
Different sets of transcripts respond to cold treatments given at different times of day in bread wheat. (A) *Triticum aestivum* cv. Cadenza seedlings were entrained under 12:12 light/dark cycles before transfer to free-running conditions. Under free running conditions, tissue was collected after 3 h cold treatments, with six such treatments given to separate batches of plants over 24 h of free running conditions. **(B)** Differentially expressed genes after a 3 h cold treatments given at six timepoints (log fold change > 0.5, FDR < 0.05). Yellow and grey shading indicate subjective day and night, respectively. **(C)** The number of differentially expressed genes (DEGs) between control temperature conditions and following a 3 h cold treatment at each timepoint. **(D)** The proportion of DEGs that only responded to a cold treatment at a single timepoint. **(E)** Summary of the difference between mean normalised control and cold transcript abundance of DEGs across the time course. Orange and teal colours indicate normalized change in cold-induced and cold-repressed transcripts, respectively.

The number of differentially expressed genes (DEGs) in response to the cold treatment was different at each at each timepoint (log fold change > 0.5, false discovery rate < 0.05; Fig. 1B). This indicates that the time of subjective day of the cold treatment determines the cold- induced transcriptome in wheat (Fig. 1B). The greatest number of genes were differentially expressed in response to a cold treatment at zeitgeber time (ZT) 48 (ZT48), with 4350 and 3403 genes significantly up or down-regulated, respectively (Fig. 1C; ZT refers to the time relative dawn, which is ZT0). Two-thirds fewer genes were differentially expressed in response to a cold treatment at ZT28 or ZT44 (Fig. 1C), despite a similar total number of transcripts detected at these timepoints (Table S1). There was a large difference in the direction of change of DEGs that responded to a cold treatment at ZT36 and ZT40, with the number of up-regulated genes comparable to those at ZT32 and ZT48, but a smaller number of down-regulated genes that was comparable to the number at ZT28 and ZT44 (Fig. 1C). These data suggest the wheat transcriptome is more responsive to a cold treatment around the middle of the subjective day (ZT32), and towards the end of the subjective night (ZT48) (Fig. 1C).

Fewer transcripts were downregulated than upregulated by an acute cold treatment given at any timepoint tested, whereas the proportion of cold-downregulated genes that were unique to each timepoint was consistently greater than the proportion upregulated genes that were unique to each timepoint (Fig. 1D). This suggests that genes downregulated in response to cold are especially influenced by the time of day. Comparing the dynamics of a representative transcript (*TaLHY*) using RNA sequencing analysis (Fig. S1A-C) and RT- qPCR (Fig. S1D) provides confidence in our transcriptomic analysis.

### The nature and cold-sensitivity of cold-responsive genes fluctuates across the 24 h cycle

We wished to understand whether the composition of the cold-responsive transcriptome changes over the 24 h cycle. To achieve this, we compared the identity of the genes that responded to a cold treatment given at each timepoint tested. Four of the largest six groups of DEGs were cold-induced at specific timepoints, so more genes responded to cold at specific (unique) times than were cold responsive at all timepoints (Fig. S2A). Furthermore, the top four sets of transcripts that were downregulated by cold were unique to single timepoints (Fig. S2B). Therefore, under free running conditions, the set of genes that responds to an acute cold treatment depends upon the time of the cold treatment. Some transcripts were only responsive to a cold treatment given at one timepoint, some across part of the 24 h cycle, and some were cold-responsive regardless of the time of the treatment (Fig. S2A, B).

Because the set of cold responsive genes changes according to the time of the cold treatment (Fig. 1D; Fig. S2A, B), we reasoned that there might be circadian gating of the responses to cold of the wheat transcriptome. There were a variety of responses to the time course of cold treatments, with cluster analysis revealing a large proportion of cold- responsive genes to have temporally-fluctuating cold response profiles (Fig. 1E). For example, certain genes are upregulated by a cold treatment given during the subjective day and down-regulated by a cold treatment given during the subjective night (Groups 5 and 6; Fig. 1E), whilst other genes are downregulated by a cold treatment during the subjective day, and upregulated or have little response to a cold treatment during the subjective night (Group 3; Fig. 1E). This indicates that the magnitude of transcriptional responses to cold in wheat can vary across a 24 hour cycle, suggesting that the circadian clock gates the transcriptional response of a variety of genes to an acute cold treatment.

### The responsiveness of signalling genes can be gated by the circadian clock

To assess how the time of day affects cold sensitivity of key signalling components, four genes that are known to be circadian regulated either in Arabidopsis or wheat were studied as examples. The hexaploid nature of the bread wheat genome means that many transcripts are present as triads, with gene homoeologs on the A, B and D subgenomes. Therefore, all three homoeologs were examined, to obtain a representative view of how a signalling component responds to a stimulus.

C-repeat binding factors (CBFs) are well-described cold responsive transcription factors [54, 55], with up to 25 CBFs present in wheat [56]. Certain *CBF* transcripts have a circadian rhythm in barley and wheat [43, 57], and in Arabidopsis the cold-induction of *CBFs* is gated by the circadian clock [28]. Our data indicate that circadian gating of *CBF* cold-responsiveness is conserved in wheat and can extend across homoeolog triads, such as for *TaCBFIVc-B14*, which is a cold-responsive *CBF* ortholog with high homology to *AtCBF2* (Fig. 2A, B, C) [24, 56]. In our experiments, all three homoeologs of *TaCBFIVc-B14* were upregulated most strongly by a cold treatment given at the end of the subjective day (Fig. 2A, B, C).

**Figure 2.**
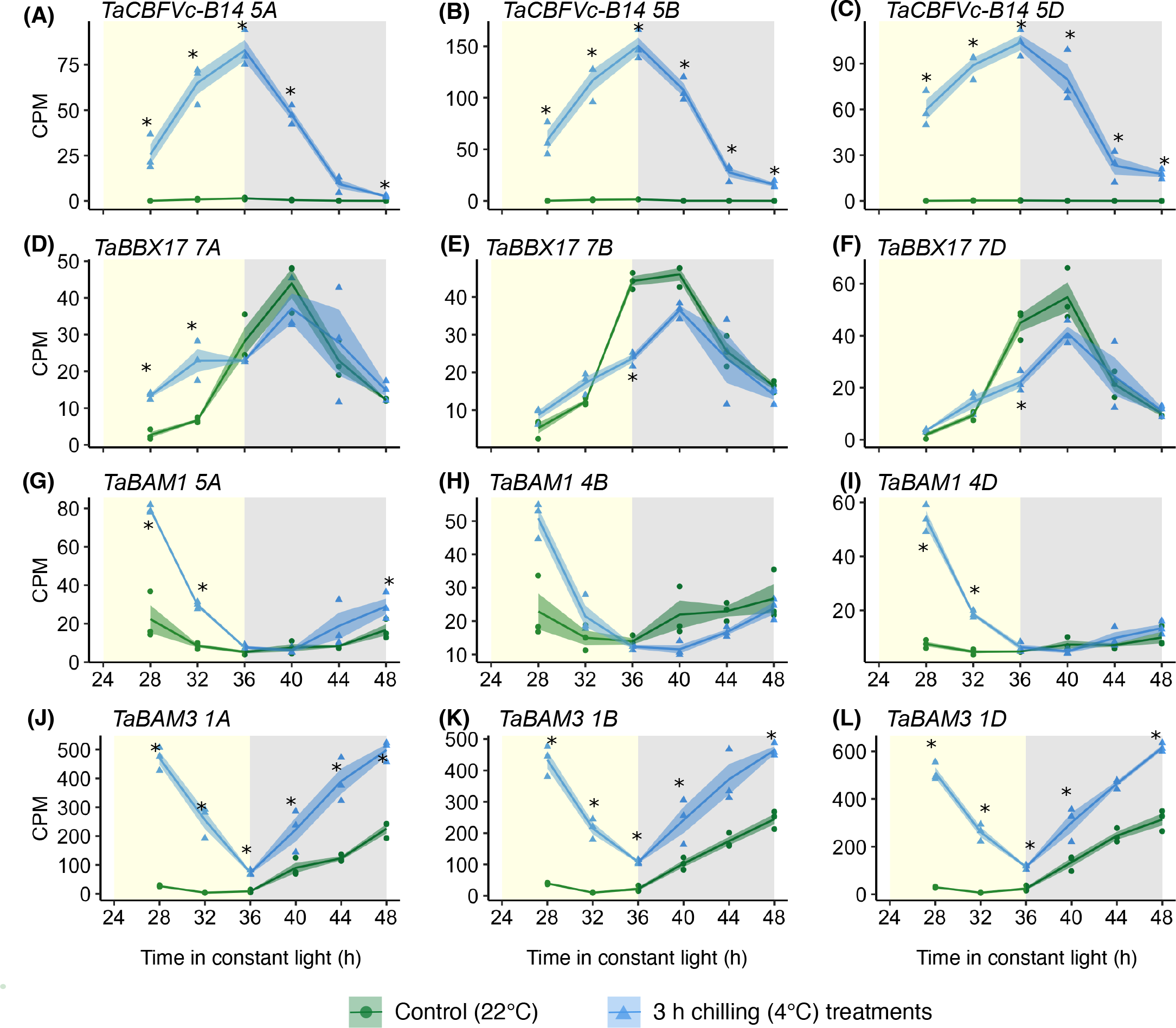
Circadian gating of the cold response of selected wheat triads. (A-C) *TaCBFVc-B14*; triad no. 15598. **(D-F)** *TaBBX17*; triad no 9634. **(G-I)** *TaBAM1*; triad no 15540. **(J-L)** *TaBAM3*; triad no 15539 [49]. Yellow/grey shading = subjective day/night. Solid lines are mean (N = 3 biological replicates; blue/green shading = ± s.e.m). Asterisks indicate times when the transcript was differentially expressed between the control and cold temperature conditions (LogFC >0.5, FDR < 0.05). Each cold treatment lasted 3 h, and ended at the time shown on the x axis. CPM (counts per million) determined by edgeR [84].

The B-BOX CONTAINING PROTEINs (BBXs) participate in responses to environmental cues [58], such as the circadian modulation of *AtBBX17* transcript responses to heat [34, 41]. Transcripts from the *TaBBX17 7ABD* triad, which has closest homology to *AtBBX17*, share with Arabidopsis rhythmic accumulation under control temperature conditions (Fig. 2D, E, F). The response to cold varied across the triad, with *TaBBX17 7A* upregulated by a cold treatment given at the start of the subjective day, but not a cold treatment given during the subjective night (Fig. 2D). This contrasts *7B* and *7D*, which were downregulated by a cold treatment given at the end of the subjective day (Fig. 2D, E, F). This example suggests that features of circadian gating are not always conserved across triads.

In Arabidopsis, the circadian clock regulates photosynthesis and starch metabolism, with these thought to provide a selective advantage to plants [59–61]. A range of related Arabidopsis transcripts encoding starch degradation enzymes are circadian regulated, but this is not always conserved in wheat, where certain orthologs of β-amylase (BAM, e.g. *BAM3*) have a circadian rhythm, whereas *BAM1* does not [43]. Under our conditions, *TaBAM1* triad and *TaBAM3* triad were upregulated by cold treatments given during the subjective day (Fig. 2G-L). *TaBAM1* was not responsive to cold treatments given during the subjective night, whereas *TaBAM3* was (Fig. 2G-L). The cold-induced relative abundance of *TaBAM3 1B* and *TaBAM3 1D* was approximately ten-fold greater than the corresponding homoeologs of *TaBAM1* (Fig. 2H, I, K, L), supporting the notion of a role for BAM3 in cold conditions.

### Circadian gating of transcriptional responses to cold is extensive, and not restricted to specific times of day

We considered circadian gating to be defined as a 24 h fluctuation in the sensitivity of a transcript to a stimulus, such that the magnitude of response to an identical stimulus given at a range of times of day oscillates with a 24 h cycle. Given that our data identified that different subsets of transcripts are respond to cold treatments given at different times of day (Fig. 1B, D, E), we reasoned that there might be widespread circadian gating of the cold-responsive transcriptome. To investigate this, we obtained a measure of the cold-sensitivity of each transcript, in response to a cold treatment given at each timepoint, by calculating the difference between the mean CPM of the transcript at control temperature and after each cold temperature treatment (Fig. 3A). This provided a measure of the response to cold, at each timepoint, that we termed ΔCPM (Fig. 3A). Using this measure, a 24 h fluctuation in the sensitivity of any given transcript to the time-series of cold treatments will manifest as an oscillation in ΔCPM (Fig. 3A, lower panel). We tested whether there was a circadian rhythm in this cold-sensitivity profile (meta2d analysis for rhythmicity; threshold of P < 0.05) (Wu *et al.*, 2016). Circadian rhythms of cold sensitivity (ΔCPM) were detected for 4388 transcripts, which we filtered to include only transcripts that were differentially expressed in response to cold at least once across the timecourse, and exclude transcripts that could not be confidently assigned to a particular time of maximum cold-responsiveness. This yielded 1677 transcripts that we classify as having circadian gating of their responses to cold in wheat (Fig. 3B).

**Figure 3.**
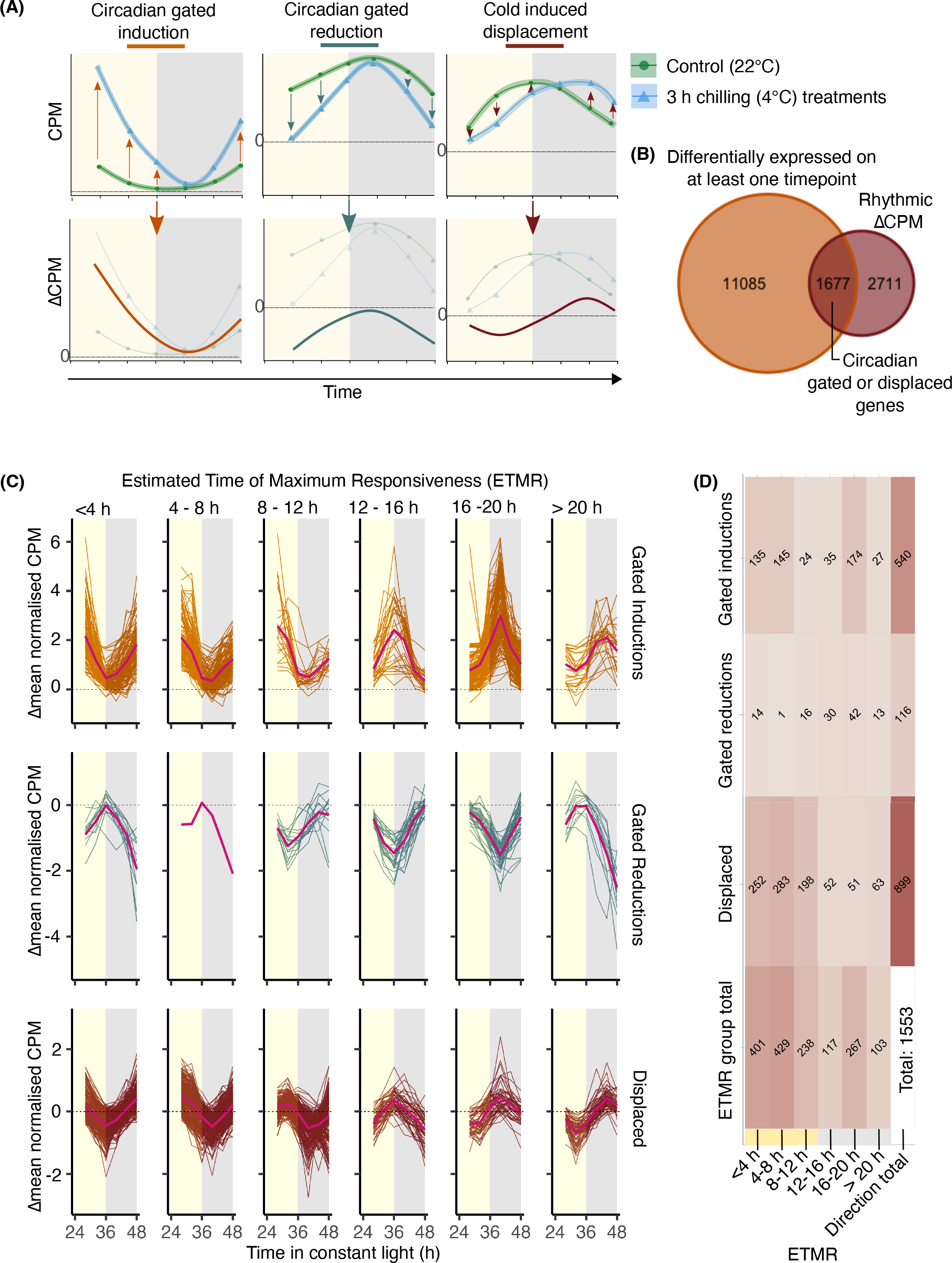
The temporal structure of the cold-responsive transcriptome. (A) We defined circadian gating of the response of genes to cold as a 24 h cycle of sensitivity of the response of the gene to identical acute (3 h) cold treatments. To identify such genes, we calculated the difference between the mean control and cold-treated CPM (termed ΔCPM). Circadian gating of a cold response manifests as a 24 h oscillation of ΔCPM, which captures cold induction, repression, or a displacement in transcript cycling. **(B)** The number of cold- responsive differentially-expressed gene transcripts (DEGs; LogFC >0.5, FDR <0.05) that have a rhythmic ΔCPM profile across the time course (meta2d, p <0.05). This represents a set of wheat transcripts with circadian gating of their response to a 3 h cold treatment. **(C)** Transcripts with circadian gating of their response to a 3 h cold treatment, organized according to their estimated time of maximum cold responsiveness (ETMR). Direction of response (induction, reduction, displacement) was based on the average difference between mean normalised control and cold-treated CPM to normalize differences in ΔCPM profile amplitude across the transcriptome. The direction was formalized as circadian-gated inductions (Δmean.normCPM > 0.5), cold induced displacement (-0.5 < Δmean.normCPM < 0.5) and circadian gated reductions (Δmean.normCPM > -0.5). ETMR was derived from meta2d phase estimates. Yellow/grey shading = subjective day/night. Pink lines indicate mean Δmean.normCPM in each directional ETMR group. **(D)** The number of genes within each directional cold ETMR group. 1533 genes were included, after filtering to remove ΔCPM profiles from ETMR groups < 4 h and > 20 h having peak CPM at an opposing timepoint from the ETMR. Darker colours indicate greater numbers of transcripts.

We found that this set of transcripts divides into three groups: circadian gated cold- inductions (where mean[ΔnormalisedCPM] > 0.5), circadian gated cold-reductions (mean[ΔnormalisedCPM] < -0.5), and a dynamic group where the transcript abundance is relatively stable for the duration of the cold treatment (mean[ΔnormalisedCPM] between -0.5 and 0.5) (Fig. 3A). Within these cold response types, we calculated the estimated time of maximum cold responsiveness (ETMR) using phase estimates from meta2d (Fig. 3C). This approach allowed us to estimate ETMR that fell between sampling timepoints, thereby overcoming some limitations of the sampling frequency. For circadian gated reductions (Fig. 3A), the time of maximum cold-responsiveness was the trough of the oscillation.

The greatest number of transcripts with rhythmic sensitivity to acute cold treatments were those whose abundance was stabilized for the duration of the cold treatment (cold-induced displacement; Fig. 3C, D). Furthermore, over 500 transcripts had circadian gating of their induction by cold (Fig. 3C, D). Within these transcripts, the most common ETMR was between 16 h and 20 h (Fig. 3D), with a large proportion of the transcripts also phasing between 4 h and 8 h. The largest proportion of cold-repressed transcripts had greatest cold-sensitivity when the cold treatment was given during the subjective night (Fig. 3D).

This indicates that there is transcriptome-wide circadian gating of the cold-responsive transcriptome in bread wheat. These responses are not restricted to a particular direction of change (upregulation *vs*. downregulation), and different transcripts have maximum responsiveness to cold treatments given at different times of day. Our analysis might under- estimate the number of circadian gated cold-responsive genes, because a longer and higher resolution time course would provide greater statistical power.

### Circadian gating of cold responses has a subtle subgenome bias

The wheat subgenomes arose from hybridisation of three diploid ancestors, resulting in a circadian oscillator that likely incorporates more components than diploid species. No specific subgenome is favoured across the circadian regulated transcriptome of wheat [43], whereas circadian regulated paralogs have differential expression patterns in other crops [39]. Furthermore, environmentally-responsive gene triads do not always have expression that is balanced across the subgenomes [49]. Therefore, we wished to determine whether there was subgenome bias in the circadian gating of transcriptomic responses to cold.

For transcripts classified as having a circadian gated cold-induction or a displaced pattern of cycling, the relative proportion of transcripts originating from A, B and D subgenomes was determined for each ETMR group (Fig. 4A). The proportion derived from each subgenome remained relatively constant in ETMR groups representing the subjective day (30 – 40%; Fig. 4A). However, during the subjective night, the largest proportion of circadian gated cold induced transcripts derived from the D subgenome (ETMR > 12 h) (Fig. 4A). This was restricted to cold-induced transcripts, and did not occur for transcripts with displaced cycling. The number of genes with circadian-gated repression by cold was rather small, so the proportional subgenome contribution to cold-repressed transcripts is noisy (Fig. S3).

**Figure 4.**
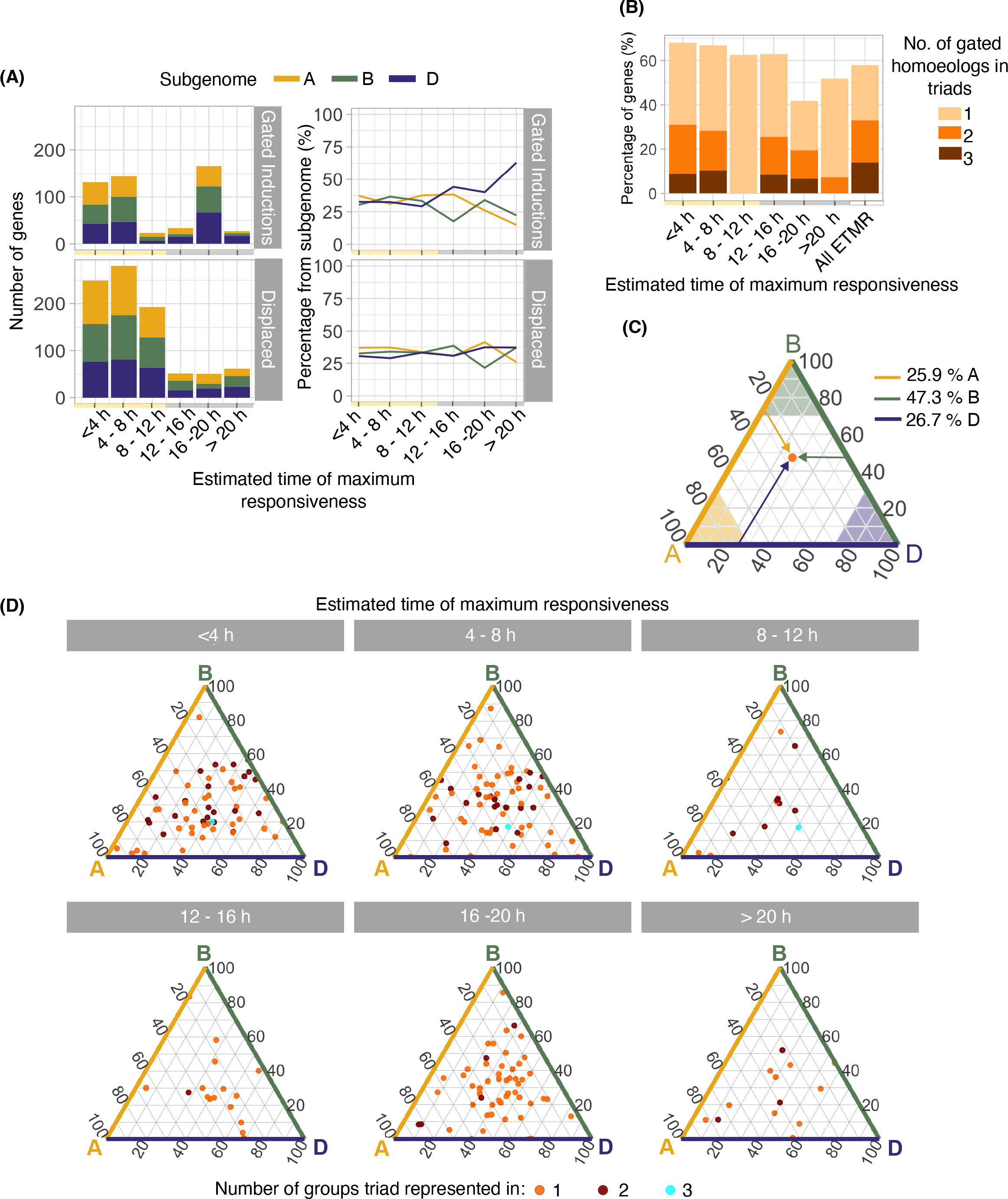
Subgenome organization of the circadian gating of the cold-responsive transcriptome. (A) The number and proportion of transcripts from each subgenome that have a circadian gated response to an acute (3 h) cold treatment, divided according to the time of maximum cold responsiveness. (**B)** The proportion of transcripts that have a circadian gated response to an acute cold treatment and belong to an annotated triad [49], divided according to the time of maximum cold responsiveness. Not all homoeologs within any given triad are gated or present within the same maximum responsiveness group, and this is indicated by the bar colour. (**C)** Example ternary plot showing the relative contribution of each homoeolog within a triad to the combined transcript accumulation from the triad [49, 101]. The position of each point on the plot reflects the proportion of total contribution from the three homoeologs within the triad (subgenome A, yellow axis; subgenome B; green axis, subgenome D, blue axis). A point in the centre of the plot represents a triad where all three homoeologs are equally cold-responsive. Points towards corners (shaded areas) represent triads where the majority of the cold response derives from a specific homoeolog. (**D)** The relative contributions of homoeologs to the cold responsiveness of each triad, organized according to the time of maximum cold responsiveness. Each triad contains at least one homoeolog that is within the ETMR group. Therefore, certain triads are present in only one ETMR (orange points), some in two (brown) and some in three (cyan).

Over 60% of circadian-gated cold induced transcripts with a ETMR before 16 h can be assigned to a triad, with a smaller proportion for transcripts with ETMR over 16 h (Fig. 4B). Approximately 10 % of transcripts with ETMR < 4 h, 4-8 h, 12-16 h and 16-20 h were detected as complete triads of three homoeologs, but no complete triads were present in the ETMR groups 12-16 h and > 20 h (Fig. 4B). Only single transcripts from triads had circadian gated responses to cold in the 8-12 h ETMR group, suggesting differential behaviour within triads (Fig. 4B). Furthermore, it appears that the time of maximum responsiveness of some triads is spread across multiple phases, with different homoeologs responding most strongly to cold treatments given at different times because the proportion of transcripts detected as complete triads across all the ETMR groups was greater than within any individual ETMR group (Fig. 4B). This suggests there is sometimes circadian gating bias within triads, with only certain homoeologs responding to the cold stimulus given at particular times of day.

The cold-responsiveness of each homoeolog to the overall cold responsiveness of each triad was assessed [49]. The relative contribution of each subgenome to the cold-responsiveness of the triad determined the position of the triad within a ternary plot (example in Fig. 4C). In general, no subgenome obviously contributed more or less than the other subgenomes (Fig. 4D). Many triads had dynamic responses to cold, in that the contribution from each subgenome to total transcript level was different under control temperature conditions compared with after the cold treatment (Fig. S4). The variation in the contribution of each subgenome to triad cold responsiveness, in each timing group, suggests that there is no overall bias towards a particular subgenome despite individual gene homoeologs occupying different EMTR groups.

### Genes associated with light responses can be circadian gated to the subjective day

Gene ontology-term analysis identified multiple biological processes associated with responses to light as over-represented in transcripts that had greatest induction by cold treatments given during the subjective day (ETMR 0-12 h). Processes such as “cellular responses to blue light”, “cellular responses to UV-A” and “anthocyanin containing compound biosynthesis” were enriched in multiple subjective day ETMR groups (Fig. S5; adjusted p < 0.001). Enrichment of terms such as “photoprotection” and “stress-activated MAPK cascade” suggests the cold stimulus elicited a broad response when delivered in the subjective day, at a time when light and low temperatures may occur simultaneously.

There was a clear split between the GO-terms that were enriched in subjective day ETMR groups and subjective night ETMR groups. “Cold acclimation” is shared across two subjective night groups (Fig. S5; adjusted p < 0.001), suggesting that night-time low temperatures could have a key role in adapting wheat to cold seasonal weather. Also enriched are terms associated with DNA transcription, which can be an indication of preparation for a large shift in gene expression (Fig. S5; adjusted p < 0.001). “Photosystem II” associated processes were enriched in subjective night ETMR group 16 – 20 h, which might relate to unexpected exposure to continuous light conditions (Fig. S5; adjusted p < 0.001). Whilst GO-term analysis interpretation requires caution, it is possible that restriction of the cold response of certain processes to specific times of day provides an advantage to wheat by ensuring that energy demanding cold-responsive processes are not induced at inappropriate times of day [62].

### Acute cold treatments displace the oscillation of certain circadian clock components

We noticed that a variety of transcripts appeared to be maintained at a relatively uniform level for the duration of the cold treatment. Under these circumstances, the transcript abundance after the acute cold treatment was similar to the cold-temperature abundance of the transcript at the previous timepoint (see example in Fig. 5A). Although this appears to be a delay in the cycling of the transcript, it does not represent a phase shift because the cycling of transcripts was not monitored following each cold treatment. Instead, this process appears to be a stabilization of the transcript abundance for the duration of the cold treatment.

**Figure 5.**
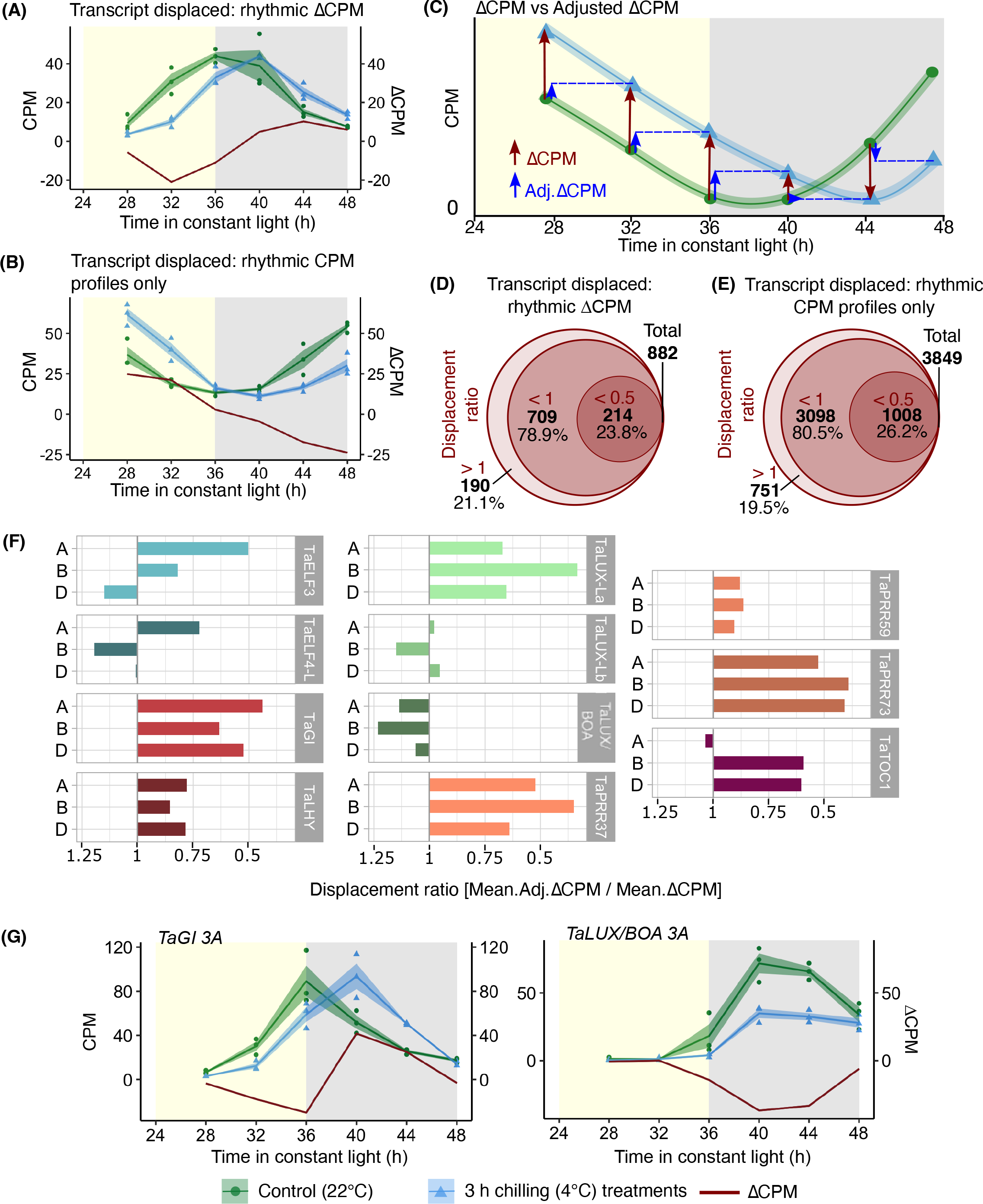
Cold temperature displaces the oscillation of a set of rhythmic transcripts. **(A)** Dynamics of a transcript with abundance stabilized during the cold treatment (*TraesCS4A02G091700*), whereby the transcript abundance after an acute cold treatment (blue) is similar to the abundance before cold treatment (green). The difference between cold and control treated transcript abundance (ΔCPM) is shown in dark red. **(B)** Dynamics of a rhythmic transcript (*TraesCS7A02G416300*; meta3d: ARS p <0.05) that has an abundance that is stabilized during the cold treatment, but has a more linear ΔCPM profile due to its underlying phase profile. **(C)** Artificial data exemplifying these dynamics. ΔCPM is the difference between the control and cold-treated CPM at each timepoint (dark red arrows), and adjusted ΔCPM is the difference between the control temperature CPM at one timepoint and the cold-treated CPM at the next timepoint (blue arrows). **(D)** A displacement ratio was calculated from the average adjusted ΔCPM as a proportion of the original average ΔCPM for the transcript, and **(E)** all transcripts with rhythmic control and cold-treated CPM profiles (meta3d: ARS p <0.05). For more details on the calculation of the displacement ratio, see Fig. S6. In **(D, E)** the larger circle represents 100% of these transcripts, the middle circle represents transcripts with a displacement ratio <1, and the smaller circle transcripts with a displacement ratio < 0.5. Displacement ratios < 1 suggest transcript abundance was stabilized for the cold treatment duration. **(F)** Displacement ratios of eleven circadian clock gene triads [43] suggest their abundance was stabilized for cold treatment duration for the majority of these transcripts. (**G)** Transcript dynamics of *TaGI 3A* (*TraesCS3A02G116300*), which had stabilized abundance during the cold treatment, and *TaLUX/BOA 3A* (*TraesCS3A02G526600*) where this did not occur. Solid lines are mean (N = 3). Blue/green shading = ± s.e.m. Red line represents ΔCPM. Yellow/grey shading = subjective day/night.

We wished to systematically identify the set of transcripts that had this response. We developed an analytical method that used the ΔCPM to consider the relationship between the transcript levels at adjacent timepoints (Fig. 5C; Fig. S6; see methods for calculations). This involved calculating an average adjusted ΔCPM for each transcript, by finding the difference between control CPM values and the adjacent cold CPM value in the timecourse. This adjusted ΔCPM value was expressed as a proportion of the original average ΔCPM of the timecourse to give a “displacement ratio” value (Fig. 5C; Fig. S6; see methods for calculations). Over three-quarters of the 882 transcripts with rhythmic ΔCPM have a displacement ratio of < 1, supporting the interpretation that the abundance of these transcripts was stabilized during the acute cold treatment (Fig. 5D). A similar proportion of the 3849 transcripts with rhythmic control and cold CPM profiles had a displacement ratio of < 1, suggesting that the abundance of these transcripts was stabilized during the cold treatment also (Fig. 5E). A possible explanation is that the cold treatment maintained a relatively constant level of these transcripts for the duration of the treatment. 47% of transcripts with circadian gated cold-induction responses had displacement ratios of < 1, suggesting around half of the transcripts with a gated response to cold did not undergo this displacement of cycling during the cold treatment.

The GO-term “circadian rhythm” was enriched within the genes that had a displacement ratio < 0.5 in response to cold, in combination with either rhythmic CPM profiles or rhythmic ΔCPM (adjusted p < 0.001; Fig. S7). This suggests that for the duration of the cold treatment, the abundance of certain circadian oscillator components was stabilized, causing an apparent displacement in the oscillation of the transcript (Fig. S6). All homoeologs of six triads of circadian clock genes [43], including morning phased *TaLHY* and evening phased *TaGI*, had a displacement ratio < 1 (Fig. 5F). Therefore, the cold treatment displaced the oscillation in the expression of these triads (Fig. 5F). Homoeologs in the triads *TaELF3*, *TaELF4-*L and *TaTOC1* had a mixture of displacement ratios, indicating that the cold treatment did not equally affect the cycling of homoeologs of all circadian clock transcripts (Fig. 5F).

Cold treatments did not have a consistent impact upon all circadian oscillator components. For example, cold had little effect upon expression of the *TaLUX-Lb* and *TaLUX/BOA* triads, which had lower or similar mean ΔCPM before adjustment for all three homoeologs (Fig. 5F). Therefore, cold does not appear to affect all circadian clock transcripts equally, as exemplified by *TaGI* having a stabilized and displaced abundance after each cold treatment, whereas *TaLUX/BOA* did not (Fig. 5G). This supports the idea that cold has an unbalanced effect upon circadian clock components, reflecting a complex relationship between the circadian clock and cold.

## Discussion

### A dual effect of cold on both the clock and clock-controlled genes

Here, we have identified for the first time extensive genome-wide circadian gating of the response to cold temperature in bread wheat. This demonstrates that the circadian gating of responses to cold of single genes [28] extends across the entire genome, and supports reports of the circadian gating of specific genes to high temperature stress in Arabidopsis [32, 40]. Given the considerable phylogenetic distance between wheat and Arabidopsis, we reason that the circadian gating of low temperature responses might be conserved across the angiosperms.

Our results demonstrate the importance of circadian regulation in shaping transcriptomic responses to short-term cold. The diversity of temporal responses to cold suggests considerable complexity in the integration of low temperature and circadian signals (Fig. 1E, Fig. 3). The presence of a circadian rhythm for a transcript under control temperature conditions does not mean that its cold-responsiveness is gated by the circadian clock, because some circadian regulated transcripts are not cold-responsive (Fig. 6A; Fig. S8A). Likewise, some transcripts are cold-responsive, but this is independent from circadian regulation because these transcripts are cold-induced by a similar magnitude, irrespective of the time of day of the cold treatment (Fig. 6B; Fig. S8B). In one form of circadian gating, the circadian clock restricts the response to the stimulus to certain times of day, and the transcript is completely unresponsive to cold treatments given at other times (Fig. 6C, Fig. S8C). This type of circadian gating is bidirectional, because suppression of transcript levels by an acute cold treatment can also be circadian-gated (Fig. 3C). In another form of circadian gating, a circadian modulation of sensitivity occurs across the day, such that a response to cold always occurs, but the response is greater in response to a cold treatment given at certain times relative to other times (e.g. *TaBAM3*; Fig. 3J-L). The phase of maximum cold responsiveness of some transcripts is sometimes aligned with the phase of the underlying circadian rhythm under control temperature conditions, but in other cases (e.g. *TaBAM3*) the phase was misaligned (Fig. 3J-L; Fig. 6D; Fig. S8D).

**Figure 6.**
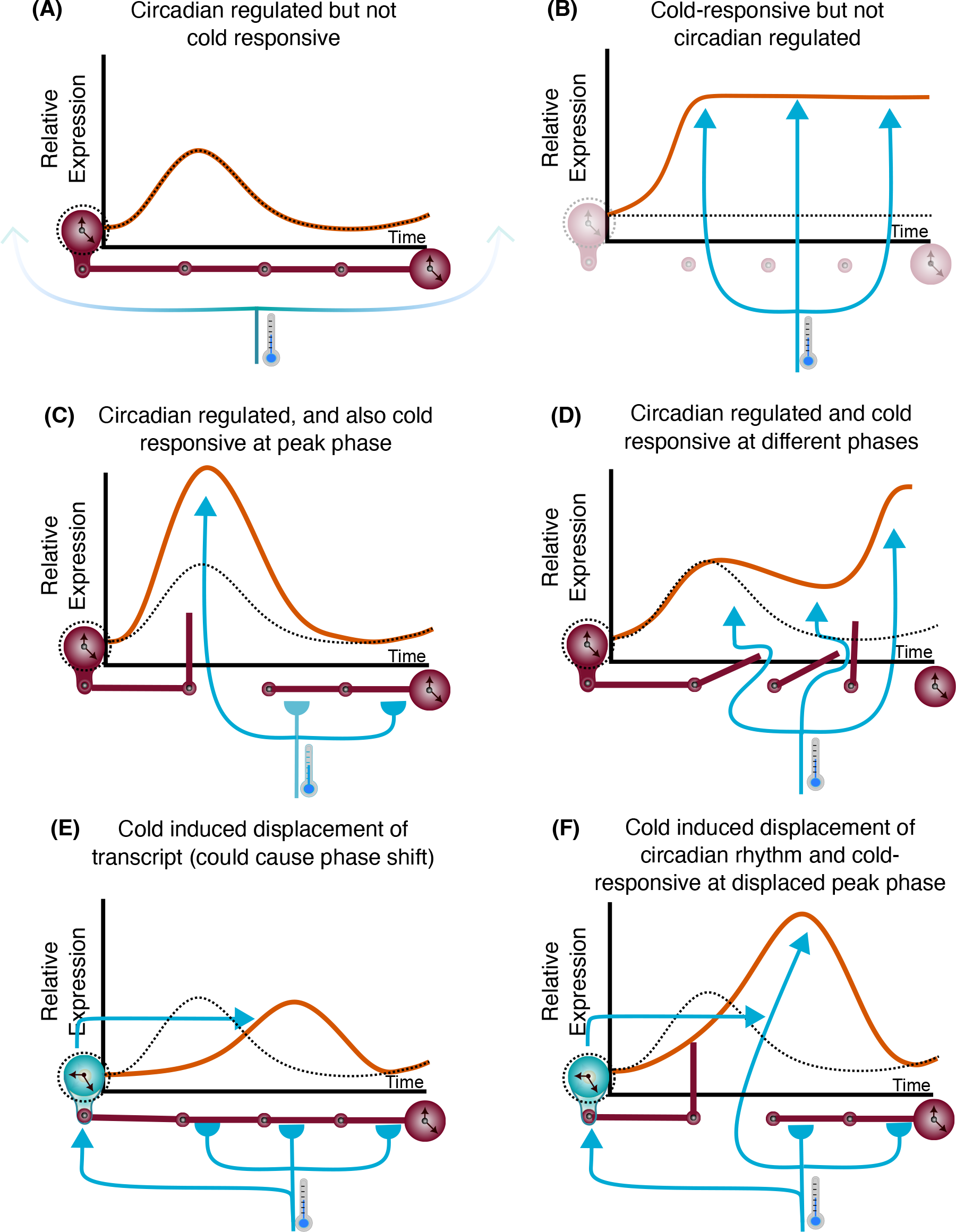
Features of the circadian gating of cold temperature responses. The interaction between the circadian clock and an acute cold treatment produces a diversity of transcript response profiles across the wheat transcriptome. Diagrams conceptualize how the circadian clock (red clocks) and cold (blue arrows) can affect underlying control- temperature transcript levels (dotted black line), producing a diversity of cold-responsive transcript profiles (orange line). **(A)** certain transcripts can be regulated by the circadian clock but transcript levels are unaltered by cold (e.g. *TraesCS5D02G447500* and *TraesCS1D02G372000*). **(B)** transcripts can respond equally to cold, irrespective of the time that the cold treatment is applied (e.g. *TraesCS2D02G284400* and *TraesCS3B02G287400*). **(C)** the cold-responsiveness of transcripts can be gated by the circadian clock, resulting in a response to an acute cold treatment at specific times of day, but not other times (e.g. *TraesCS3D02G325300)*. **(D)** the circadian clock gates the response to cold of the transcript, with the time of maximum responsiveness to cold having a different phase from the phase of the underlying circadian rhythm of the transcript (e.g. *TaBAM3*). **(E)** cold affects the circadian oscillator directly (cold blue clock), stabilizing oscillator transcript levels for the duration of the acute cold treatment (e.g. *TaGI* 3A, *TraesCS5B02G271300*). **(F)** the cold- responsiveness of a transcript is restricted to a particular phase, and cold also displaces the oscillation of the circadian clock transcript, apparently by stabilizing its abundance for the duration of the cold treatment. In combination, this upregulates and delays the accumulation of the transcript (e.g. *TraesCS1A02G289800* and *TraesCS1B02G095000*).

Short cold treatments affect the dynamics of certain wheat circadian clock components, by stabilising their abundance for approximately the duration of the cold treatment (Fig. 6E). This pervades hundreds of other circadian-regulated cold-responsive transcripts (Fig. 6F). It is possible that this would cause a cold-induced delay in the circadian oscillator under free running conditions, leading to comparable delays in the response of downstream circadian-regulated transcripts. Interestingly, this occurs for some circadian clock genes (e.g. *TaPRR73* and *TaGI*) but not others (e.g. *LUX* orthologues) (Fig. 5F). One potential explanation for this bifurcating response is that the stability of certain transcripts is insensitive to cold, so the transcript level continues to change under low temperature conditions. Another explanation could be that different cell types in the leaves of wheat have differential temperature responses which, combined with a predominance of different circadian clock components in different cell types [63], causes bifurcated dynamics of distinct circadian clock components in response to cold. Uneven temperature responses have been reported across the components of the Arabidopsis circadian oscillator [64], so a similar situation might occur in wheat. Finally, an open and speculative question- for which there is not currently evidence- is of whether polyploid genomes, such as the genome of hexaploid wheat, can ever support partial decoupling of subgenome-derived circadian oscillators in single cells. It will in future be informative to know whether this cold-induced stabilization of oscillator transcript levels can act as a zeitgeber for the wheat circadian clock, in a manner reminiscent of the phase shifts caused in Arabidopsis by moderate temperature reductions [65]. This is important because it could contribute to the phase relationship between circadian-regulated processes and environmental phase under field conditions, with consequences for metabolism or flowering time. It will also be informative to understand conserved and differing aspects of these responses in winter wheat, given its different vernalization and photoperiod requirements compared with spring wheat varieties such as Cadenza, used here. Taken together, these results suggest that fluctuations in environmental temperature might cause a dynamic adjustment of circadian phase in cultivated wheat [66].

Our data suggest two broad processes underlying the dynamics of circadian regulated and cold-responsive transcripts in spring wheat. In the first case, low temperature stabilizes the abundance of circadian oscillator transcripts. It would be informative in future to determine whether this shifts the phase of the circadian oscillator. In the second case, the circadian oscillator constrains or amplifies the influence upon a gene of a cold treatment given at certain times, compared with other times. This oscillation in cold-sensitivity can have a phase that is similar to or different from the phase of the underlying rhythm of expression of the transcript. This is clearly demonstrated by the cold responsiveness of *TaBAM3*, for which the phase at which a cold treatment gives the greatest response is not defined by the phase of the transcript under control temperature conditions (Fig. 2J-L). This suggests that for some genes, separate circadian signalling components might regulate underlying rhythms and cold responsiveness.

### Potential cold induction bias towards D subgenome during subjective night

Genes that had greatest responsiveness to a cold treatment during the subjective day had a balanced contribution of transcripts from each subgenome. In contrast, for transcripts with largest induction by a cold treatment given during the subjective night, the greatest proportion of transcript derived from the D subgenome (Fig. 4A). However, when comparing the relative contributions from homoeologs that are present as triads, there did not seem to be a bias towards the D subgenome (Fig. 4E). Therefore, this greater proportion of transcripts from the D subgenome could arise from genes that are present as diads or singletons. This might relate to the origins of the D subgenome donor, *Aegilops tauschii*. The D subgenome hybridizations are thought to involve *A. tauschii* derived from mountainous regions near the Caspian Sea [67, 68], so perhaps the greater cold induction from the D subgenome during the subjective night relates to adaptation of *A. tauschii* to relatively cold conditions.

### Relationship between the circadian gating of the cold-responsive transcriptome and circadian clock outputs

Rhythms in transcript abundance do not always lead to rhythms in protein abundance or activity [69–72], although there is circadian gating of the high temperature-responsive translatome [41]. In many cases, the abundance of proteins encoded by rhythmic transcripts remains constant [72–74], whilst in some cases (e.g. BAM3 in Arabidopsis) the protein abundance tracks the transcript abundance [74]. Circadian regulation of protein turnover occurs in eukaryotes [75, 76], and it is thought that circadian regulation of gene expression might maintain proteostasis in the presence of rhythmic protein degradation [77]. This might explain why rhythms in protein abundance are less prevalent than rhythms of transcript abundance.

This idea could be relevant to interpreting the physiological significance of circadian gating in wheat. For approximately half of the genes that have a circadian gated response to cold, the phase of maximum cold sensitivity was similar to the phase of the rhythm at control temperatures (Fig. 6C). This alignment of the phase of maximum cold sensitivity with the phase of the underlying rhythm likely increases the availability of mRNA at certain times, perhaps in anticipation of a greater rate of protein turnover [78]. This idea is exemplified by the greatest sensitivity to cold treatments given during the subjective day of genes that have light signalling GO terms (Fig. S4). We reason that this temporal regulation increases transcript levels for these components, in anticipation of cold- and light-induced oxidative damage in the daytime. Conversely, transcripts that have circadian gating of their downregulation by cold might encode proteins that are stabilised in cold- so do not need to be replaced- or contribute to processes that are not preferred in cold conditions. Alternatively, down regulated genes may represent repressive regulators, like many components of the circadian clock [79], so a reduction in their abundance could lift repression of cold response mechanisms.

## Conclusions

We identified that the circadian clock plays a key role in shaping the cold responsive transcriptome through the process of circadian gating. This establishes that circadian regulation is central to the temperature responses of a major cereal crop. Our study supports the notion that circadian or temporal gating of responses to environmental cues represents a widespread regulatory mechanism in plants, from peach trees and grape vines [80, 81] to Arabidopsis [34], and here at a genome-wide scale in hexaploid bread wheat. Our findings open opportunities to understand the contributions of circadian regulation to wheat crop performance, and could underpin biotechnological developments that optimize the resilience of cereals to an increasingly unpredictable climate.

## Materials and methods

### Plant materials and growth conditions

Seeds of *Triticum aestivum* cv. Cadenza (donated by James Simmonds, John Innes Centre) were stratified on damp filter paper for 3 days at 4°C before germination at 22°C in darkness for two days. Seedlings were cultivated in 24-cell trays for 12 days on a bespoke cereal cultivation mixture (65% peat, 25% loam, 10% horticultural grit, 3 kg m^−3^ dolomitic limestone, 1.3 kg m^−3^ PG Mix fertiliser (Yara), 3 kg m^−3^ osmocote extract) under cycles of 12 h light and 12 h darkness, at approximately 200 μmol m^−2^ s^−1^ of white light at 22°C (Fig. S9A).

### Circadian time course sampling

14-day old wheat seedlings at Zadok stage GS1.2 were transferred to constant light for 24 h before experimentation commenced, to ensure data were free from transitory effects caused by the final dawn (Fig. S9B). After 25 h of constant conditions (zeitgeber time (ZT) 25, where ZT 0 refers to the start of the day), three individuals were transferred to 4°C with identical light conditions for 3 h, whilst control plants remained at 22 °C. After the 3 h cold treatment, 3 cm of tissue from the second leaf was collected from each control and treated plant (indicated by red boxes on Fig. S9B), immediately frozen in liquid N_2_, and stored at -80°C until processing. After sampling, the cold-treated and control plants were not used for subsequent samples. Six such further treatments were conducted, beginning at ZT 25, ZT 29, ZT 33, ZT 37, ZT 41 and ZT 45, and ending at ZT 28, ZT 32, ZT 36, ZT 40, ZT 44 and ZT 48, respectively (Fig. 1A). Each set of samples used a different batch of plants.

### RNA sequencing data collection, QC and read mapping

RNA was extracted using Macherey-Nagel NucleoSpin RNA Mini Kit with on-column DNase treatment, according to manufacturer’s instructions. RNA was stored at -80°C in nuclease- free water. RNA integrity was assessed using an Agilent 2100 Bioanalyzer System at the John Innes Centre, and again by our sequencing partner Novogene (Cambridge, UK). All samples had RIN scores > 6 and concentrations > 50 ng RNA μl^-1^, thereby passing quality control. Libraries were constructed by Novogene using a mRNA polyA enrichment library protocol involving fragmentation, cDNA synthesis, end repair, A-tailing, adapter ligation, size selection, amplification and purification. Libraries were verified on a Bioanalyzer before pooling for sequencing. After verification, 150bp paired-end sequencing was performed on the Illumina NovaSeq 6000 platform to generate a minimum of 15 GB data per sample, generating 675.5 GB total data with an average read depth of 62.5 million reads per sample. All samples had ≥ 98.5 % clean reads suitable for alignment.

### RNA sequencing analysis

Transcript quantification was conducted using the RNA-seq pseudoaligner kallisto v0.44.0 (Bray et al., 2016) using 31 bp k-mers and default parameters. The index file was constructed from the high confidence CDS sequence from the International Wheat Genome Sequencing Consortium (IWGSC) *T. aestivum* cv. Chinese Spring RefSeqv1.1 [48].

### Differential gene expression analysis

Kallisto transcript count data were condensed to the gene level using R v2022.07.1 [82]. The Bioconductor [83] packages edgeR [84–86] and Limma [87] were used to identify differentially expressed genes between cold-treated and control seedlings at each timepoint. Counts per million (CPM) were determined by edgeR and low-expressed transcripts filtered out of the data (edgeR function filterByExpr, which uses the filtering strategy described in Chen *et al.* 2016) (Dataset S1, Dataset S2). This filtering strategy retains genes with a CPM value above *k* in *n* samples, where *k* is determined from the set minimum count value and sample size, and *n* determined from the smallest group size within the design matrix [86]. Around 68.5% of transcripts remained after filtering (Dataset S3). Empirical Bayes moderation was performed for predictions of gene-wise variability [88] and a correction for multiple testing (Benjamini-Hochberg method) was used to obtain the false discovery rate (FDR) of differentially expressed genes. Differentially expressed genes were defined as those with an adjusted p-value < 0.05 and logFC > 0.5 (50% change).

### Data processing and visualisation

CPM and logFC values were assembled using base R and the tidyverse package dplyr [89, 90]. Graphs were produced using ggplot2 [91] and radar charts with fmsb function radarchart [92]. Upset plots (Fig. S2) were created with UpSetR [93] and GO-terms presented using code adapted from [43, 94]. The sensitivity heatmap was produced using mean normalised CPM values determined by dividing the average CPM at any given timepoint by the mean CPM for each gene across the entire time course under both temperature conditions. The heatmap was created with ComplexHeatmap, using the Ward D.2 method for clustering, with k = 12 [95, 96].

### Identification of genes having a circadian gated response to cold

We used the difference between the mean control transcript level and mean cold-treated transcript level (ΔCPM) at each timepoint as a measure of the sensitivity of gene expression to cold. To test for rhythmicity of cold sensitivity, the ΔCPM profiles were analysed across the entire timecourse duration using the MetaCycle v1.2.0 function meta2d [97] (Fig. 3A; Fig. S6). The minimum and maximum permitted period lengths were 20 h and 28 h, respectively [98]. We considered meta2d to be especially suitable for this analysis because it integrates period, phase and p-values from three algorithms, reducing the limitations of individual algorithms, and is appropriate for analysis of a relatively small number of time-points [98]. A rhythmicity cut-off of meta2d p-value < 0.05 was used. In general, time-series close to this p-value threshold had approximately 24 h rhythmicity (Fig. S10). As an additional filter, only genes with rhythmic ΔCPM that were also differentially expressed at least once in the time-series were considered to be circadian gated. Only transcripts with average CPM > 1 under control temperature conditions or cold temperature conditions, across the time-series, were included in the analysis to eliminate low-expressed genes with large relative variation between replicates due to the low counts. The difference between mean normalised CPM values (Δmean-normCPM) was used to assign transcripts to three directional response groups: circadian-gated inductions by cold (Δmean-normCPM > 0.5), cold induced displacement of transcript cycling (-0.5 < Δmean-normCPM < 0.5) and circadian gated reductions by cold (Δmean-normCPM > -0.5).

The estimated time of maximum cold responsiveness (ETMR) refers to the time of the greatest ΔCPM. This was estimated from the phase measured using meta2d analysis, which provides the peak phase of the time-series data. For the ETMR for circadian gated cold reductions, the ETMR was determined using meta2d phase estimates for the inverse ΔCPM values, because they represent the phase of the trough. This allowed a total of 18 directional ETMR groups to be defined (Fig. 3C). Extra filters were applied to ETMR groups < 4 h and > 20 h for inductions and displaced transcripts, to remove 135 trend anomalies that had greater ΔCPM at the opposite timepoint (+ 12 h) to the meta2d phase estimate.

### Gene ontology-term analysis

GO-term enrichment analysis for biological processes was performed using TopGo [99] for each cold-induction ETMR group. The group sizes were sufficiently large for this approach (n>20). GO-terms were retrieved from the IWGSC RefSeqv1.0 annotation and transferred to annotation v1.1 using the method described by [100]. Enrichment analysis was performed on gene lists through comparison against a gene universe of 75251 genes, which comprised all genes expressed in our dataset that had high quality annotation and could be associated with a GO term (nodeSize = 10, algorithm = weight01) [99]. WeightedFisher values were adjusted for multiple testing using the Benjamini-Hochberg correction method to generate adjusted p-values, filtered stringently for adjusted p < 0.001. The same pipeline was used for genes with rhythmic control and cold CPM profiles, and rhythmic ΔCPM that had a displacement of abundance for the duration of the cold treatment (displacement ratio <0.5).

### Investigation of the contribution of wheat subgenomes to triad responsiveness

Subgenome occupancy was determined using information within the gene identifier (e.g. TraesCS**<u>1A</u>**02G123456). Genes were organised into triads with a 1:1:1 correspondence using the triad numbers in the High Confidence Triad table from [49] (https://github.com/Uauy-Lab/ WheatHomoeologExpression/Data/data/TablesForExploration/HCTriads.csv). Triads were considered to be part of a circadian gated induction ETMR group if any homoeolog from the triad was categorised into the group. Individual homoeologs of triads were sometimes within different ETMR groups (Fig. 4D). An average ΔCPM value for the ETMR for each gene was determined as:

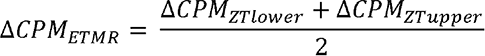

Where is the ZT timepoint corresponding to the lower time limit of the ETMR group and ZTupper is the ZT timepoint corresponding to the upper time limit of the ETMR group, e.g for ETMR group 4 – 8 h, ZTlower = ZT 28 and ZTupper = ZT 32. The relative contribution of each homoeolog to the triad responsiveness for the ETMR was determined as:

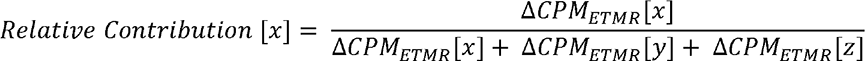

where is the subgenome of the homoeolog being investigated, and and are the remaining two subgenomes. This used a method similar to Ramirez-Gonzalez et al. [49]. The relative contributions were determined and visualised in R using ggtern [101].

### Identification of transcripts where cycling is stabilized during an acute cold treatment

We wished to identify the entire set of transcripts that had a stabilization of abundance during the cold treatments. To achieve this, transcripts with rhythmicity at control temperatures and in response to 3 h cold treatments were identified using the ARSER algorithm in Metacycle (using meta3d) [97, 102]. Genes previously defined as having a circadian gated upregulation or downregulation in response to cold were removed from this group to prevent double-counting of individual genes. Adjusted ΔCPM (Adj.ΔCPM; Fig. S6) values were determined for timepoints ZT28-44 for each gene as:

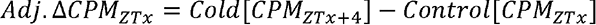

Where ZT is the is sampling timepoint. A time course mean ΔCPM for each gene was determined as:

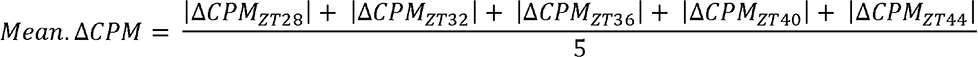

And a mean for adjusted ΔCPM for each gene was determined as:

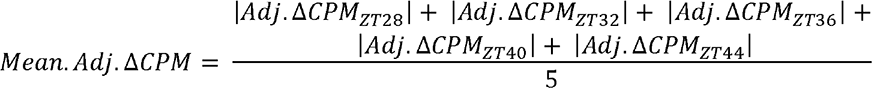

A displacement ratio value was determined for each gene in this group using mean ΔCPM and mean adjusted ΔCPM values as:

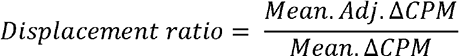

Displacement ratios less than 1 indicate a reduced mean ΔCPM following adjustment. Gene identifiers for putative wheat circadian clock genes were obtained from [43], and displacement ratios determined for each.

### cDNA synthesis and RT-qPCR analysis

For RT-qPCR experiments, the amount of RNA was standardised across samples to to 0.1 μg/μl and cDNA was synthesised using a High-Capacity cDNA Reverse Transcription Kit and random primers (Applied Biosystems). Transcript abundance in cDNA dilutions of 1/100 was measured in 25 µl reactions using qPCRBIO SyGreen LO-ROX RT-qPCR reagent (PCR Biosystems) with relevant primers on a CFX96 qPCR machine (Bio-Rad, California, US). Transcript abundance was normalized to *TaHK4* (*TraesCS5A02G015600*) [103] according to the ΔΔCT method. Primers (5’ -> 3’) were ATCAATAATGGTACTTCTCCGGG (forward) and TGATTTTGGAACTTCTCTGGTG (reverse) for *TaLHY*, and TCTAAATGTCCAGGAAGCTGTTA (forward) and CCTGTGGTGCCCAACTATT (reverse) for *TaHK4*.

## Acknowledgements

We thank James Simmonds for seeds of *T. aestivum* cv. Cadenza, Paige Panter for experimental design advice, Cristobal Uauy and Philippa Borrill for advice on data interpretation, and Hannah Rees and Rachel Rusholme-Pilcher for advice on data processing. Figure S6 was created with biorender.com.

## Funding

BBSRC Institute Strategic Programme GEN BB/P013511/1 (AND, PP).

The Leverhulme Trust RPG-2018-216 (AND).

SWBio DTP grants BB/J014400/1 and BB/M009122/1 (CAG, KJE, AND).

## Author contributions

Conceptualization: CAG, KJE, AND.

Investigation: CAG, PP.

Analysis and visualization: CAG, PP.

Writing: CAG, PP, KJE, AND.

## Data availability

Raw sequencing data are deposited as European Nucleotide Archive dataset PRJEB56923. Processed data are provided as supplemental information within the manuscript.

Figure S1. **Comparison of *TaLHY* transcript dynamics using RNA sequencing analysis and RT-qPCR.** (A-C) Relative abundance of *TaLHY* homoeologs from the A, B and D wheat subgenomes quantified using RNA sequencing analysis. (D) Relative abundance of *TaLHY* transcript abundance (using primers binding all homoeologs) using RT-qPCR analysis. Solid lines are mean (N = 3). Blue/green shading = ± s.e.m. Yellow/grey shading = subjective day/night.

Figure S2. **The number DEGs shared between timepoints.** Upset plots visualising the size of the intersections between the groups of genes **(A)** up-regulated, and **(B)** down- regulated by an acute cold treatment each timepoint. In a similar manner to Venn diagrams, filled circles signify the timepoint of interest and the links between filled circles signify the intersection of interest, the size of which is reported by the height of the bar immediately above. Set size represents the total amount of DEGs detected at each timepoint.

Figure S3. **The number and proportion of transcripts from each subgenome that have a circadian gated down regulated response to cold.** Small group sizes for each Estimated Time of Maximum Responsiveness group resulted in large variation in the proportion of transcripts derived from each subgenome.

Figure S4. **Dynamic contribution of homoeologs to triad cold responsiveness.** The relative contributions of homoeologs to total triad transcript abundance following each cold treatment and in its respective control, split by the estimated time of maximum responsiveness (ETMR) groups. Ternary plots as described in Fig. 4C and cold treatment data is that presented in Fig. 4D.

Figure S5. **Gene ontology-term analysis of transcripts that have circadian gating of their induction by cold.** GO term enrichment within each estimated time of maximum responsive cold upregulation. Circle size represents the number of transcripts associated with the GO term, and circle colour indicates the Benjamini-Hochberg adjusted weighted Fisher *p*-value.

Figure S6. **Explanation of use of the adjusted** Δ**CPM to identify transcripts having a displaced response to an acute cold treatment.** The top row of diagrams shows a hypothetical transcript that has a circadian gated response to cold, with the phase of the gate aligned with the phase of the oscillation under control temperature conditions. The lower row of diagrams shows a hypothetical transcript that has a displaced response to cold. Under these circumstances, the CPM at any given timepoint is similar to the CPM of the cold-treated sample at the subsequent timepoint, because the transcript level changes little during the cold treatment. This feature is identified from the ratio of the average ΔCPM to the average adjusted ΔCPM.

Figure S7. **Gene ontology-term analysis of transcripts that have a temporal displacement due to the cold treatment.** GO term enrichment for transcripts with a displacement ratio < 0.5 in response to cold, in combination with either rhythmic CPM profiles or rhythmic ΔCPM (meta3d p <0.05). Circle size represents the number of transcripts associated with the GO term, and circle colour indicates the Benjamini-Hochberg adjusted weighted Fisher *p*-value (adjusted p < 0.001).

Figure S8. **Transcripts exemplifying regulatory features described in** Figure 6. Transcripts **(A)** *TraesCS5D02G447500*, **(B)** *TraesCS3B02G287400*, **(C)** *TraesCS3D02G325300*, **(D)** *TraesCS5A02G468300*, **(E)** *TraesCS5B02G271300*, **(F)** *TraesCS5D02G076400*. Yellow/grey shading = subjective day/night. Solid lines are mean (N = 3 biological replicates). Blue/green shading = ± s.e.m. Red line represents ΔCPM.

Figure S9. **The light spectra used for cultivation and appearance of seedlings that were sampled**. (**A)** Light spectra used for plant cultivation and experimentation, for control and cold temperature conditions. (**B)** Representative images of bread wheat seedlings after germination on damp filter paper, followed by 12 days growth on compost under 12 h : 12 h light dark cycles (22°C) followed by 24 h of constant light. Seedlings were equivalent to Zadok Stage GS1.2 at time of sampling. Red boxes indicate region of second leaf that was sampled. Scale bar 20 mm.

Figure S10. Transcripts at the upper limit of meta2d statistical threshold for rhythmicity are candidate genes with circadian gating of their response to cold. Example transcripts that have rhythmic ΔCPM (meta2d p <0.05) and p-value close to the cut-off limit, demonstrating the level of statistical stringency was appropriate for detection of circadian-gated transcripts. Transcripts are from ETMR grouping **(A)** < 4 h, *TraesCS1A02G210400*, **(B)** 4 – 8 h, *TraesCS5D02G343300*, **(C)** 8 – 12 h, *TraesCS5D02G318100*, **(D)** 12 – 16 h, *TraesCS5A02G311100*, **(E)** 16 – 20 h, *TraesCS3B02G342936*, **(F)** > 20 h, *TraesCSU02G072000*. Yellow/grey shading = subjective day/night. Solid lines are mean (N = 3 biological replicates). Blue/green shading = ± s.e.m. Red line represents ΔCPM.

**Dataset S1.** Gene-level counts for entire time-series and transcriptome.

**Dataset S2.** Counts per million data for entire time-series and transcriptome.

**Dataset S3.** The number of genes considered from each timepoint/treatment replicate and the number of genes considered before and after Voom/Limma filtering of low-expressed genes.

## References

1. Harmer, S.L., et al., Orchestrated transcription of key pathways in Arabidopsis by the circadian clock. Science, 2000. 290(5499): p. 2110–2113.

2. Aschoff, J., Temporal orientation: circadian clocks in animals and humans. Animal Behaviour, 1989. 37: p. 881–896.

3. Millar, A.J., The intracellular dynamics of circadian clocks reach for the light of ecology and evolution. Annual Review of Plant Biology, 2016. 67: p. 595–618.

4. Covington, M.F., et al., Global transcriptome analysis reveals circadian regulation of key pathways in plant growth and development. Genome Biology, 2008. 9: p. R130.

5. Hsu, P.Y., U.K. Devisetty, and S.L. Harmer, Accurate timekeeping is controlled by a cycling activator in Arabidopsis. eLife, 2013. 2: p. e00473.

6. Kamioka, M., et al., Direct repression of evening genes by CIRCADIAN CLOCK- ASSOCIATED1 in the Arabidopsis circadian clock. The Plant Cell, 2016. 28: p. 696–711.

7. Nakamichi, N., et al., Transcriptional repressor PRR5 directly regulates clock-output pathways. Proceedings of the National Academy of Sciences, 2012. 109: p. 17123–17128.

8. Gendron, J.M., et al., Arabidopsis circadian clock protein, TOC1, is a DNA-binding transcription factor., in Proceedings of the National Academy of Sciences of the United States of America. 2012. p. 3167–3172.

9. Nakamichi, N., Adaptation to the local environment by modifications of the photoperiod response in crops. Plant and Cell Physiology, 2015. 56: p. 594–604.

10. McClung, C.R., Circadian clock components offer targets for crop domestication and improvement. Genes, 2021. 12: p. 374.

11. Oravec, M.W. and K. Greenham, The adaptive nature of the plant circadian clock in natural environments. Plant Physiology, 2022. 190: p. 968–980.

12. Steed, G., et al., Chronoculture, harnessing the circadian clock to improve crop yield and sustainability. Science, 2021. 372: p. 479.

13. Gu, L., et al., The 2007 eastern US spring freeze: Increased cold damage in a warming world? BioScience, 2008. 58: p. 253-262.

14. Trnka, M., et al., Adverse weather conditions for European wheat production will become more frequent with climate change. Nature Climate Change, 2014. 4: p. 637–643.

15. Barlow, K.M., et al., Simulating the impact of extreme heat and frost events on wheat crop production: A review. Field Crops Research, 2015. 171: p. 109–119.

16. Whaley, J.M., et al., Frost damage to winter wheat in the UK: the effect of plant population density. European Journal of Agronomy, 2004. 21: p. 105–115.

17. Thakur, P., et al., Cold stress effects on reproductive development in grain crops: An overview. Environmental and Experimental Botany, 2010. 67: p. 429–443.

18. Li, X., et al., Winter wheat photosynthesis and grain yield responses to spring freeze. Agronomy Journal, 2015. 107: p. 1002–1010.

19. Leonardos, E.D., et al., Daily photosynthetic and C-export patterns in winter wheat leaves during cold stress and acclimation. Physiologia Plantarum, 2003. 117: p. 521–531.

20. Jame, Y.W. and H.W. Cutforth, Simulating the effects of temperature and seeding depth on germination and emergence of spring wheat. Agricultural and Forest Meteorology, 2004. 124: p. 207–218.

21. Hurry, V.M. and N.P.A. Huner, Effect of cold hardening on sensitivity of winter and spring wheat leaves to short-term photoinhibition and recovery of photosynthesis. Plant Physiology, 1992. 100: p. 1283–1290.

22. Subedi, K.D., et al., Cold temperatures and boron deficiency caused grain set failure in spring wheat (Triticum aestivum L.). Field Crops Research, 1998. 57: p. 277–288.

23. Venzhik, Y.V., et al., Influence of lowered temperature on the resistance and functional activity of the photosynthetic apparatus of wheat plants. Biology Bulletin, 2011. 38: p. 132–137.

24. Winfield, M.O., et al., Plant responses to cold: transcriptome analysis of wheat. Plant Biotechnology Journal, 2010. 8: p. 749–771.

25. Fuller, M.P., et al., The freezing characteristics of wheat at ear emergence. European Journal of Agronomy, 2007. 26: p. 435–441.

26. Hotta, C.T., et al., Modulation of environmental responses of plants by circadian clocks. Plant, Cell & Environment, 2007. 30: p. 333–349.

27. Skopik, S.D. and C.S. Pittendrigh, Circadian systems, II. The oscillation in the individual Drosophila pupa; its independence of developmental stage. Proceedings of the National Academy of Sciences, 1967. 58: p. 1862–1869.

28. Fowler, S.G., D. Cook, and M.F. Thomashow, Low temperature induction of Arabidopsis CBF1, 2, and 3 is gated by the circadian clock. Plant Physiology, 2005. 137: p. 961-968.

29. Dodd, A.N., et al., Time of day modulates low-temperature Ca signals in Arabidopsis. The Plant Journal, 2006. 48: p. 962–973.

30. Bieniawska, Z., et al., Disruption of the Arabidopsis circadian clock is responsible for extensive variation in the cold-responsive transcriptome. Plant Physiology, 2008. 147: p. 263–279.

31. Zhu, J.-Y., et al., TOC1–PIF4 interaction mediates the circadian gating of thermoresponsive growth in Arabidopsis. Nature Communications, 2016. 7: p. 13692.

32. Blair, E.J., et al., Contribution of time of day and the circadian clock to the heat stress responsive transcriptome in Arabidopsis. Scientific Reports, 2019. 9: p. 4814.

33. Grinevich, D.O., et al., Novel transcriptional responses to heat revealed by turning up the heat at night. Plant Molecular Biology, 2019. 101: p. 1–19.

34. Bonnot, T., et al., Circadian coordination of cellular processes and abiotic stress responses. Current Opinion in Plant Biology, 2021. 64: p. 102133.

35. Kidokoro, S., et al., Posttranslational regulation of multiple clock-related transcription factors triggers cold-inducible gene expression in Arabidopsis. Proceedings of the National Academy of Sciences, 2021. 118: p. e2021048118.

36. Li, B., et al., Transcriptional profiling reveals a time-of-day-specific role of REVEILLE 4/8 in regulating the first wave of heat shock–induced gene expression in Arabidopsis. The Plant Cell, 2019. 31(10): p. 2353–2369.

37. Nagano, A.J., et al., Deciphering and prediction of transcriptome dynamics under fluctuating field conditions. Cell, 2012. 151: p. 1358–1369.

38. Cano-Ramirez, D.L., et al., Circadian and environmental signal transduction in a natural population of Arabidopsis. bioRxiv, 2022.

39. Greenham, K., et al., Expansion of the circadian transcriptome in Brassica rapa and genome-wide diversification of paralog expression patterns. eLife, 2020. 9: p. e58993.

40. Dickinson, P.J., et al., Chloroplast signaling gates thermotolerance in Arabidopsis. Cell Reports, 2018. 22(7): p. 1657–1665.

41. Bonnot, T. and D.H. Nagel, Time of the day prioritizes the pool of translating mRNAs in response to heat stress. The Plant Cell, 2021. 33: p. 2164–2182.

42. Calixto, C.P.G., et al., Rapid and dynamic alternative splicing impacts the Arabidopsis cold response transcriptome. The Plant Cell, 2018. 30(7): p. 1424–1444.

43. Rees, H., et al., Circadian regulation of the transcriptome in a complex polyploid crop. PLOS Biology, 2022. 20: p. 1–37.

44. Beales, J., et al., A Pseudo-Response Regulator is misexpressed in the photoperiod insensitive Ppd-D1a mutant of wheat (Triticum aestivum L.). Theoretical and Applied Genetics, 2007. 115: p. 721–733.

45. Bentley, A.R., et al., Short, natural, and extended photoperiod response in BC2F4 lines of bread wheat with different Photoperiod-1 (Ppd-1) alleles. Journal of Experimental Botany, 2013. 64: p. 1783–1793.

46. Shaw, L.M., A.S. Turner, and D.A. Laurie, The impact of photoperiod insensitive Ppd- 1a mutations on the photoperiod pathway across the three genomes of hexaploid wheat (Triticum aestivum). The Plant Journal, 2012. 71: p. 71–84.

47. Turner, A., et al., The pseudo-response regulator Ppd-H1 provides adaptation to photoperiod in Barley. Science, 2005. 310: p. 1031–1034.

48. IWGSC, Shifting the limits in wheat research and breeding using a fully annotated reference genome. Science, 2018. 361: p. eaar7191.

49. Ramírez-González, R.H., et al., The transcriptional landscape of polyploid wheat. Science, 2018. 361: p. 6089.

50. Mikkelsen, M.D. and M.F. Thomashow, A role for circadian evening elements in cold- regulated gene expression in Arabidopsis. The Plant Journal, 2009. 60: p. 328–339.

51. Dong, M.A., E.M. Farré, and M.F. Thomashow, CIRCADIAN CLOCK-ASSOCIATED 1 and LATE ELONGATED HYPOCOTYL regulate expression of the C-REPEAT BINDING FACTOR (CBF) pathway in Arabidopsis. Proceedings of the National Academy of Sciences, 2011. 108: p. 7241–7246.

52. Fowler, D.B. and A.E. Limin, Interactions among factors regulating phenological development and acclimation rate determine low-temperature tolerance in wheat. Annals of botany, 2004. 94: p. 717–724.

53. Li, Q., et al., Transcriptomic insights into phenological development and cold tolerance of wheat grown in the field. Plant Physiology, 2017. 176: p. 2376–2394.

54. Thomashow, M.F., Molecular basis of plant cold acclimation: insights gained from studying the CBF cold response pathway. Plant Physiology, 2010. 154: p. 571–577.

55. Thomashow, M.F., et al., Role of the Arabidopsis CBF transcriptional activators in cold acclimation. Physiologia Plantarum, 2001. 112: p. 171–175.

56. Badawi, M., et al., The CBF gene family in hexaploid wheat and its relationship to the phylogenetic complexity of cereal CBFs. Molecular Genetics and Genomics, 2007. 277: p. 533–554.

57. Gierczik, K., et al., Circadian and light regulated expression of CBFs and their upstream signalling genes in barley. International Journal of Molecular Sciences, 2017. 18.

58. Xu, D., COP1 and BBXs-HY5-mediated light signal transduction in plants. New Phytologist, 2020. 228: p. 1748–1753.

59. Graf, A., et al., Circadian control of carbohydrate availability for growth in Arabidopsis plants at night. Proceedings of the National Academy of Sciences, 2010. 107: p. 9458–9463.

60. Smith, S.M., et al., Diurnal changes in the transcriptome encoding enzymes of starch metabolism provide evidence for both transcriptional and posttranscriptional regulation of starch metabolism in Arabidopsis leaves. Plant Physiology, 2004. 136: p. 2687–2699.

61. Dodd, A.N., et al., Plant circadian clocks increase photosynthesis, growth, survival, and competitive advantage. Science, 2005. 309: p. 630–633.

62. Markham, K.K. and K. Greenham, Abiotic stress through time. New Phytologist, 2021. 231: p. 40–46.

63. Endo, M., et al., Tissue-specific clocks in Arabidopsis show asymmetric coupling. Nature, 2014. 515: p. 419–422.

64. Michael, T.P., P.A. Salomé, and C.R. McClung, Two Arabidopsis circadian oscillators can be distinguished by differential temperature sensitivity. Proceedings of the National Academy of Sciences, 2003. 100: p. 6878–6883.

65. Salomé, P.A. and C.R. McClung, What makes the Arabidopsis clock tick on time? A review on entrainment. Plant, Cell & Environment, 2005. 28: p. 21–38.

66. Webb, A.A.R., et al., Continuous dynamic adjustment of the plant circadian oscillator. Nature Communications, 2019. 10: p. 550.

67. Gaurav, K., et al., Population genomic analysis of Aegilops tauschii identifies targets for bread wheat improvement. Nature Biotechnology, 2022. 40: p. 422–431.

68. Zhou, Y., et al., Triticum population sequencing provides insights into wheat adaptation. Nature Genetics, 2020. 52: p. 1412–1422.

69. Baerenfaller, K., et al., Systems-based analysis of Arabidopsis leaf growth reveals adaptation to water deficit. Molecular Systems Biology, 2012. 8: p. 606.

70. Choudhary, M.K., et al., Quantitative circadian phosphoproteomic analysis of arabidopsis reveals extensive clock control of key components in physiological, metabolic, and signaling pathways. Molecular & Cellular Proteomics, 2015. 14: p. 2243–2260.

71. Graf, A., et al., Parallel analysis of Arabidopsis circadian clock mutants reveals different scales of transcriptome and proteome regulation. Open Biology, 2017. 7: p. 160333.

72. Seaton, D.D., et al., Photoperiodic control of the Arabidopsis proteome reveals a translational coincidence mechanism. Molecular Systems Biology, 2018. 14: p. e7962.

73. Uhrig, R.G., et al., Diurnal dynamics of the Arabidopsis rosette proteome and phosphoproteome. Plant, Cell & Environment, 2021. 44: p. 821–841.

74. Krahmer, J., et al., The circadian clock gene circuit controls protein and phosphoprotein rhythms in Arabidopsis thaliana. Molecular & Cellular Proteomics, 2022. 21.

75. Li, L., et al., Protein degradation rate in Arabidopsis thaliana leaf growth and development. The Plant Cell, 2017. 29: p. 207–228.

76. Srikanta, S.B. and N. Cermakian, To Ub or not to Ub: Regulation of circadian clocks by ubiquitination and deubiquitination. Journal of Neurochemistry, 2021. 157: p. 11–30.

77. Seinkmane, E., et al., Circadian regulation of protein turnover and proteome renewal. bioRxiv, 2022.

78. Murata, N., et al., Photoinhibition of photosystem II under environmental stress. Biochimica et Biophysica Acta, 2007. 1767: p. 414–421.

79. Fogelmark, K. and C. Troein, Rethinking transcriptional activation in the arabidopsis circadian clock. PLOS Computational Biology, 2014. 10: p. 1–12.

80. Artlip, T.S., et al., CBF gene expression in peach leaf and bark tissues is gated by a circadian clock. Tree Physiology, 2013. 33: p. 866–877.

81. Rienth, M., et al., Day and night heat stress trigger different transcriptomic responses in green and ripening grapevine (vitis vinifera) fruit. BMC Plant Biology, 2014. 14: p. 108.

82. Team, R.C., R: A language and environment for statistical computing. 2018, R Foundation for Statistical Computing, Vienna, Austria.

83. Huber, W., et al., Orchestrating high-throughput genomic analysis with Bioconductor. Nature Methods, 2015. 12: p. 115–121.

84. Robinson, M.D., D.J. McCarthy, and G.K. Smyth, edgeR: a Bioconductor package for differential expression analysis of digital gene expression data. Bioinformatics, 2010. 26: p. 139–140.

85. McCarthy, D.J., Y. Chen, and G.K. Smyth, Differential expression analysis of multifactor RNA-Seq experiments with respect to biological variation. Nucleic Acids Research, 2012. 40: p. 4288–4297.

86. Chen, Y., A.A.T. Lun, and G.K. Smyth, From reads to genes to pathways: differential expression analysis of RNA-Seq experiments using Rsubread and the edgeR quasi- likelihood pipeline. F1000Research, 2016. 5: p. 1438.

87. Ritchie, M.E., et al., limma powers differential expression analyses for RNA- sequencing and microarray studies. Nucleic Acids Research, 2015. 43: p. e47.

88. Smyth, G.K., Linear models and empirical bayes methods for assessing differential expression in microarray experiments. Statistical Applications in Genetics and Molecular Biology, 2004. 3: p. Article 3.

89. Wickham, H., et al., Welcome to the tidyverse. Journal of Open Source Software, 2019. 4: p. 1686.

90. Wickham, H., et al., dplyr: A grammar of data manipulation. 2022.

91. Wickham, H., ggplot2: Elegant graphics for data analysis. 2016.

92. Nakazawa, M., fmsb: functions for medical statistics book with some demographic data. 2022.

93. Gehlenborg, N., UpSetR: a more scalable alternative to Venn and Euler Diagrams for visualizing intersecting sets. 2019.

94. De Vega, J.J., et al., Differential expression of starch and sucrose metabolic genes linked to varying biomass yield in Miscanthus hybrids. Biotechnology for Biofuels, 2021. 14: p. 98.

95. Gu, Z., Complex heatmap visualization. iMeta, 2022.

96. Gu, Z., R. Eils, and M. Schlesner, Complex heatmaps reveal patterns and correlations in multidimensional genomic data. Bioinformatics, 2016.

97. Wu, G., et al., MetaCycle: an integrated R package to evaluate periodicity in large scale data. Bioinformatics, 2016. 32: p. 3351–3353.

98. Wu, G., et al., MetaCycle: Evaluate periodicity in large scale data. 2019.

99. Alexa, A. and J. Rahnenfuhrer, topGO: Enrichment Analysis for Gene Ontology. 2022.

100. Borrill, P., et al., Identification of transcription factors regulating senescence in wheat through gene regulatory network modelling. Plant Physiology, 2019. 180: p. 1740–1755.

101. Hamilton, N.E. and M. Ferry, *ggtern: Ternary Diagrams Using ggplot2.* Journal of Statistical Software, Code Snippets, 2018. 87: p. 1–17.

102. Yang, R. and Z. Su, Analyzing circadian expression data by harmonic regression based on autoregressive spectral estimation. Bioinformatics, 2010. 26: p. i168–i174.

103. Borrill, P., R. Ramirez-Gonzalez, and C. Uauy, expVIP: a customizable RNA-seq data analysis and visualization platform Plant Physiology, 2016. 170(4): p. 2172–2186.

