## Supplemental Figures S1-S10 for "Genome-wide circadian gating of a cold temperature response in bread wheat"

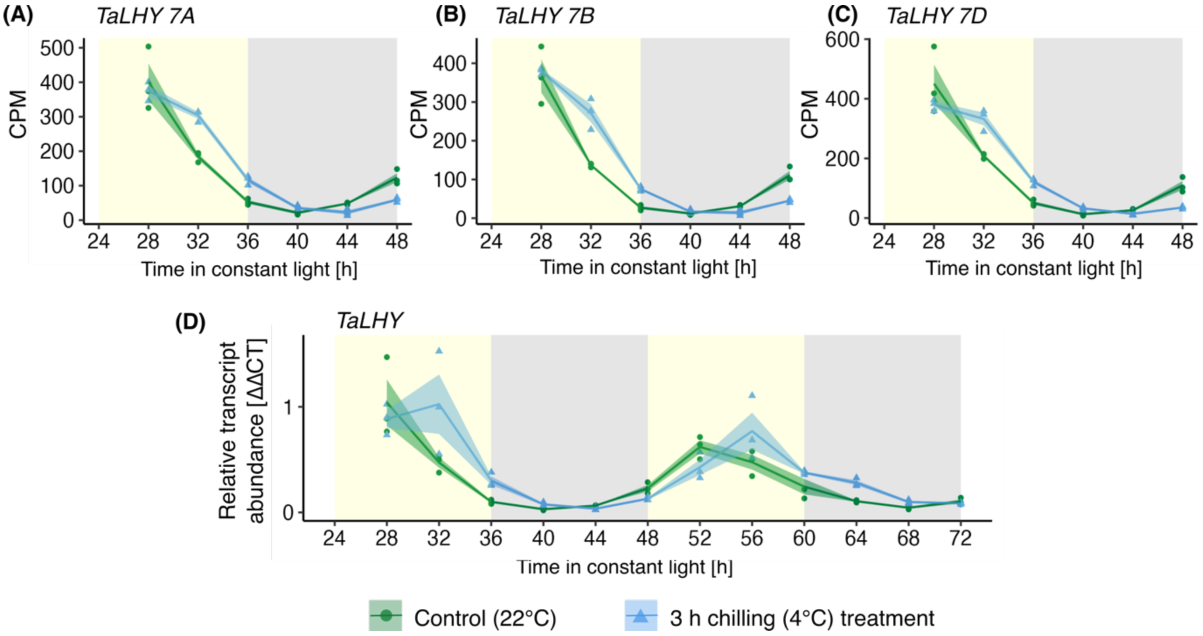


**Figure S1. Comparison of *TaLHY* transcript dynamics using RNA sequencing analysis and RT-qPCR.** (A-C) Relative abundance of *TaLHY* homoeologs from the A, B and D wheat subgenomes quantified using RNA sequencing analysis. (D) Relative abundance of *TaLHY* transcript abundance (using primers binding all homoeologs) using RT-qPCR analysis. Solid lines are mean (N = 3). Blue/green shading = ± s.e.m. Yellow/grey shading = subjective day/night.

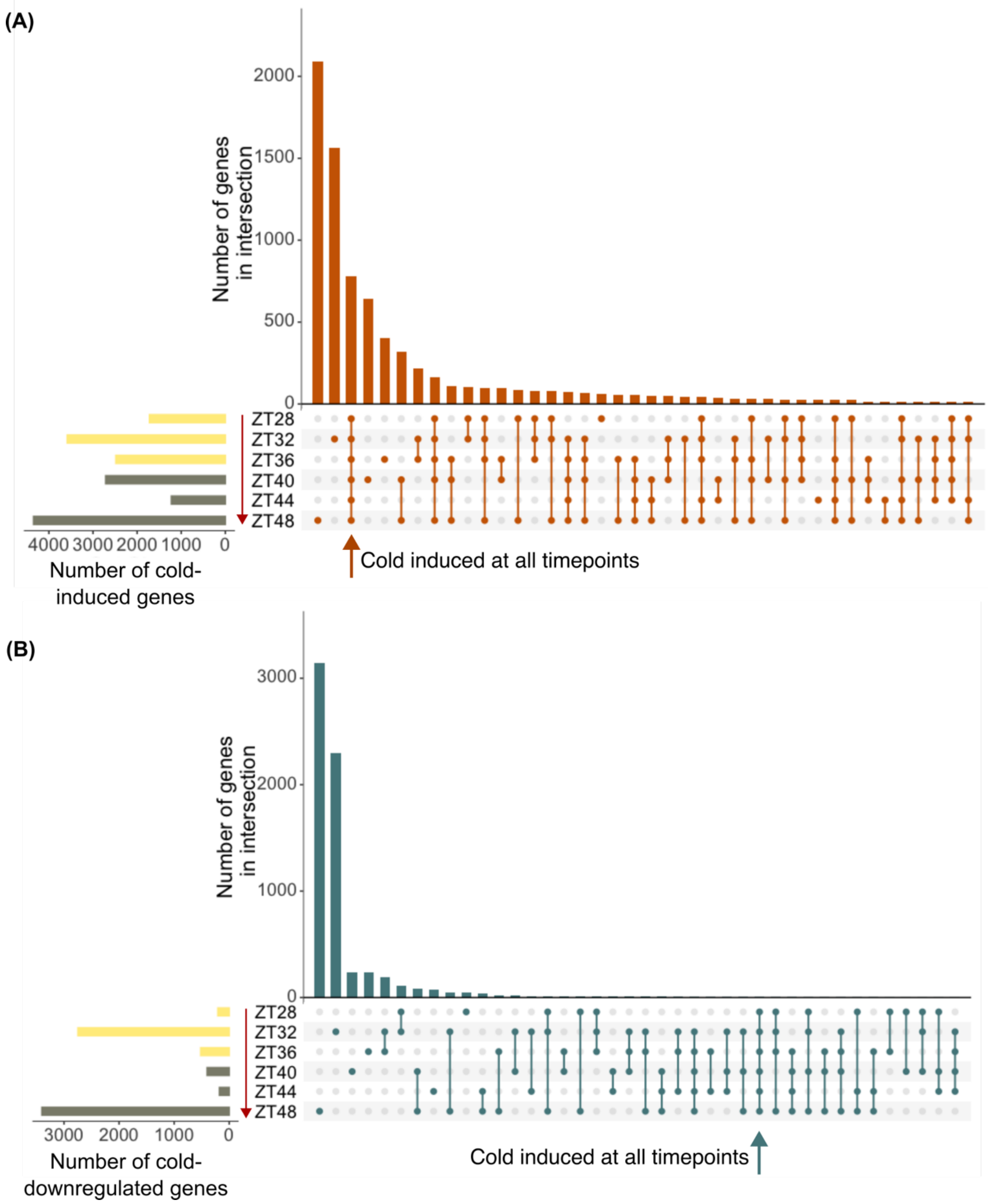


**Figure S2. The number DEGs shared between timepoints.** Upset plots visualising the size of the intersections between the groups of genes **(A)** up-regulated, and **(B)** down-regulated by an acute cold treatment each timepoint. In a similar manner to Venn diagrams, filled circles signify the timepoint of interest and the links between filled circles signify the intersection of interest, the size of which is reported by the height of the bar immediately above. Set size represents the total amount of DEGs detected at each timepoint.

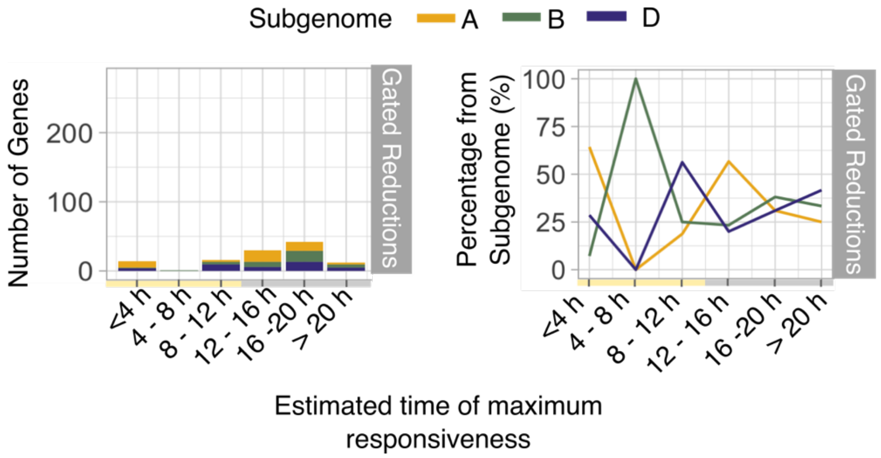


**Figure S3. The number and proportion of transcripts from each subgenome that have a circadian gated down regulated response to cold.** Small group sizes for each Estimated Time of Maximum Responsiveness group resulted in large variation in the proportion of transcripts derived from each subgenome.

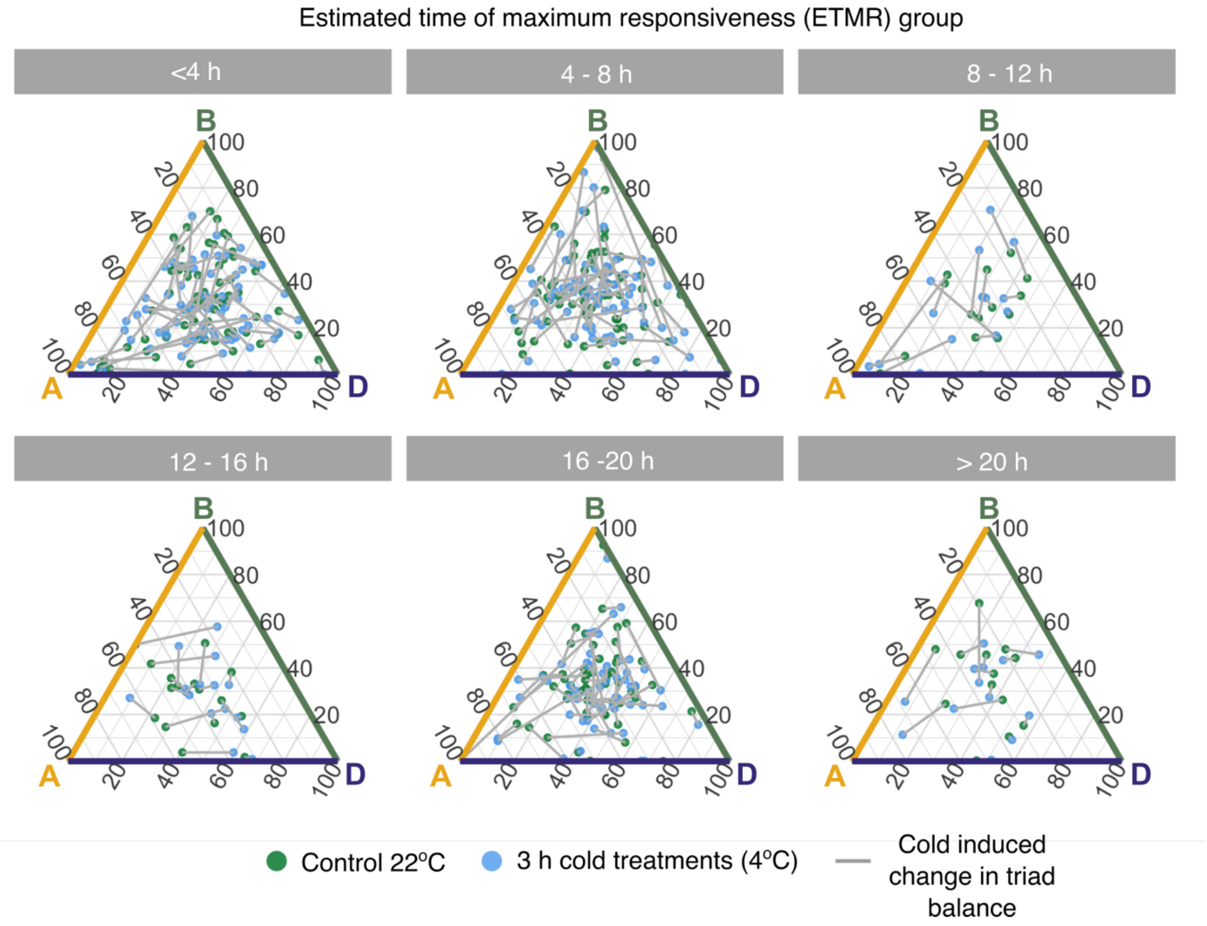


**Figure S4. Dynamic contribution of homoeologs to triad cold responsiveness.** The relative contributions of homoeologs to total triad transcript abundance following each cold treatment and in its respective control, split by the estimated time of maximum responsiveness (ETMR) groups. Ternary plots as described in Fig. 4C and cold treatment data is that presented in Fig. 4D.

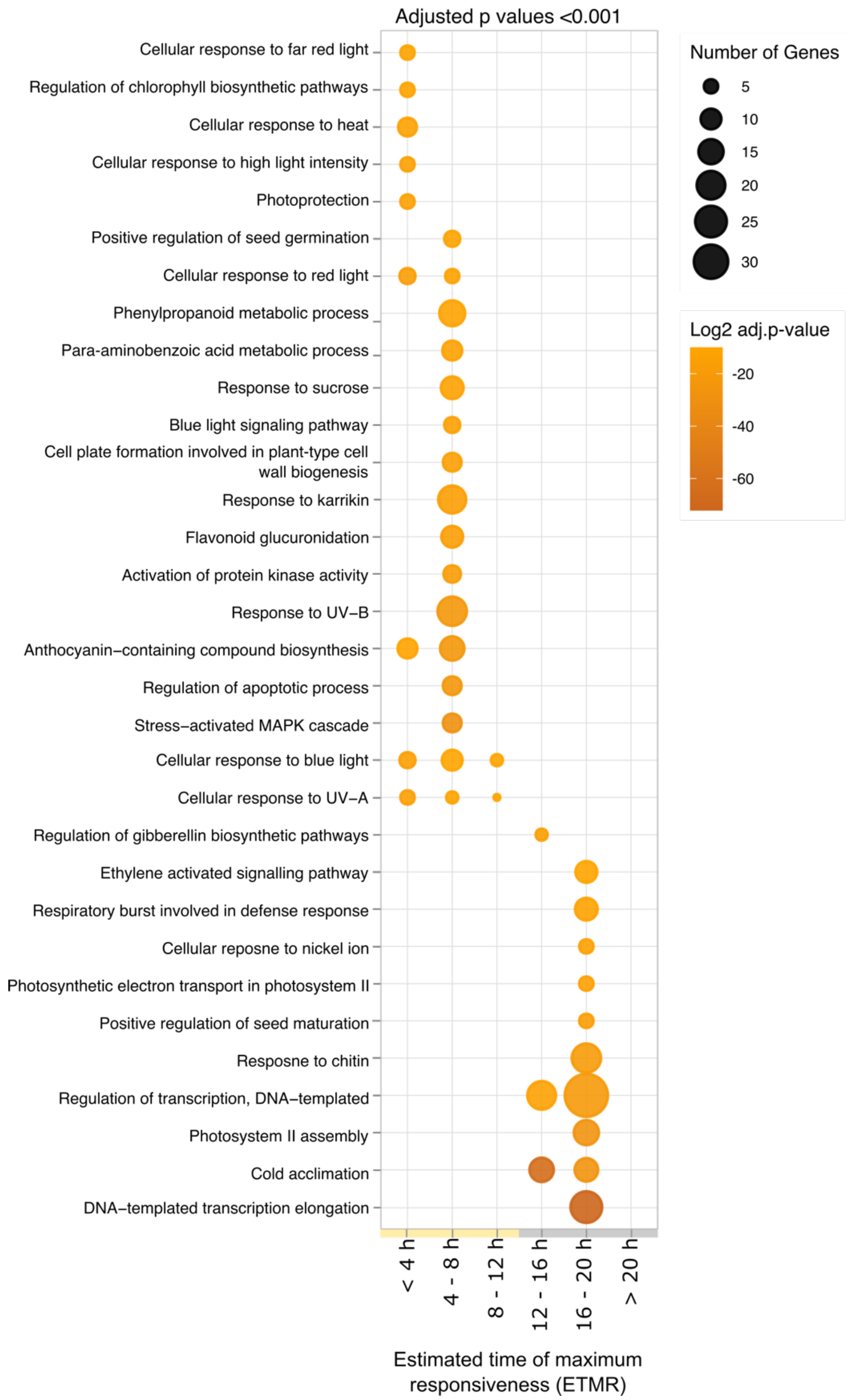


**Figure S5. Gene ontology-term analysis of transcripts that have circadian gating of their induction by cold.** GO term enrichment within each estimated time of maximum responsive cold upregulation. Circle size represents the number of transcripts associated with the GO term, and circle colour indicates the Benjamini-Hochberg adjusted weighted Fisher *p*-value.

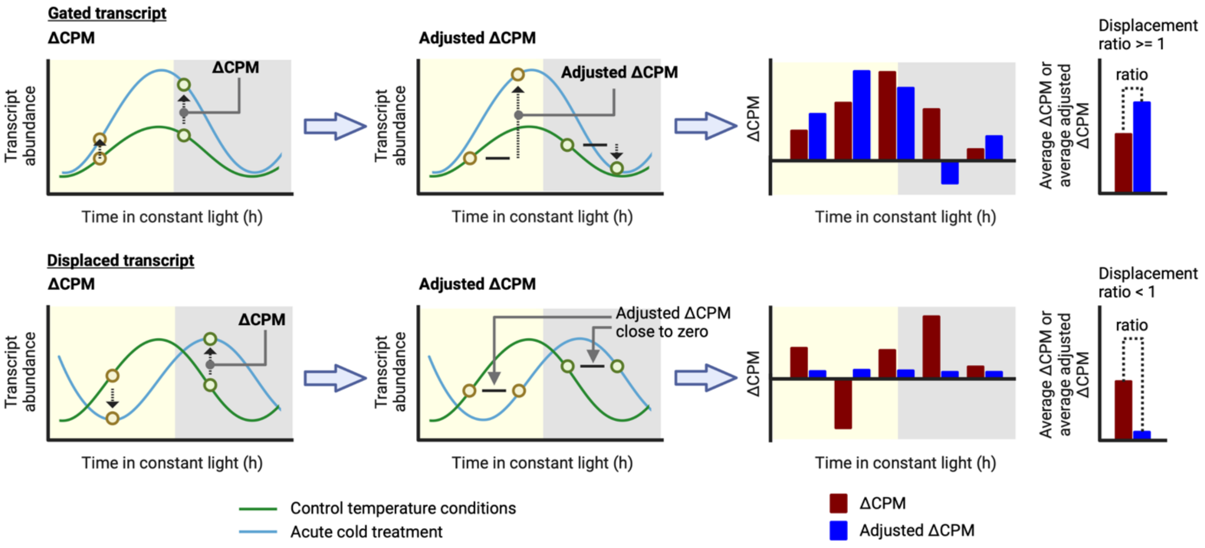


**Figure S6. Explanation of use of the adjusted ∆CPM to identify transcripts having a displaced response to an acute cold treatment.** The top row of diagrams shows a hypothetical transcript that has a circadian gated response to cold, with the phase of the gate aligned with the phase of the oscillation under control temperature conditions. The lower row of diagrams shows a hypothetical transcript that has a displaced response to cold. Under these circumstances, the CPM at any given timepoint is similar to the CPM of the cold-treated sample at the subsequent timepoint, because the transcript level changes little during the cold treatment. This feature is identified from the ratio of the average ∆CPM to the average adjusted ∆CPM.

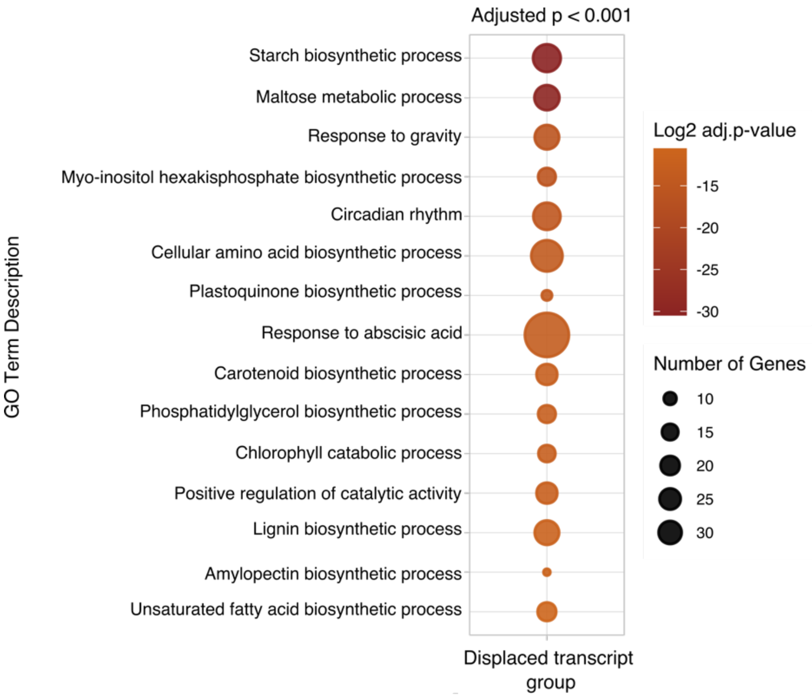


**Figure S7. Gene ontology-term analysis of transcripts that have a temporal displacement due to the cold treatment.** GO term enrichment for transcripts with a displacement ratio < 0.5 in response to cold, in combination with either rhythmic CPM profiles or rhythmic ∆CPM (meta3d p <0.05). Circle size represents the number of transcripts associated with the GO term, and circle colour indicates the Benjamini-Hochberg adjusted weighted Fisher *p*-value (adjusted p < 0.001).

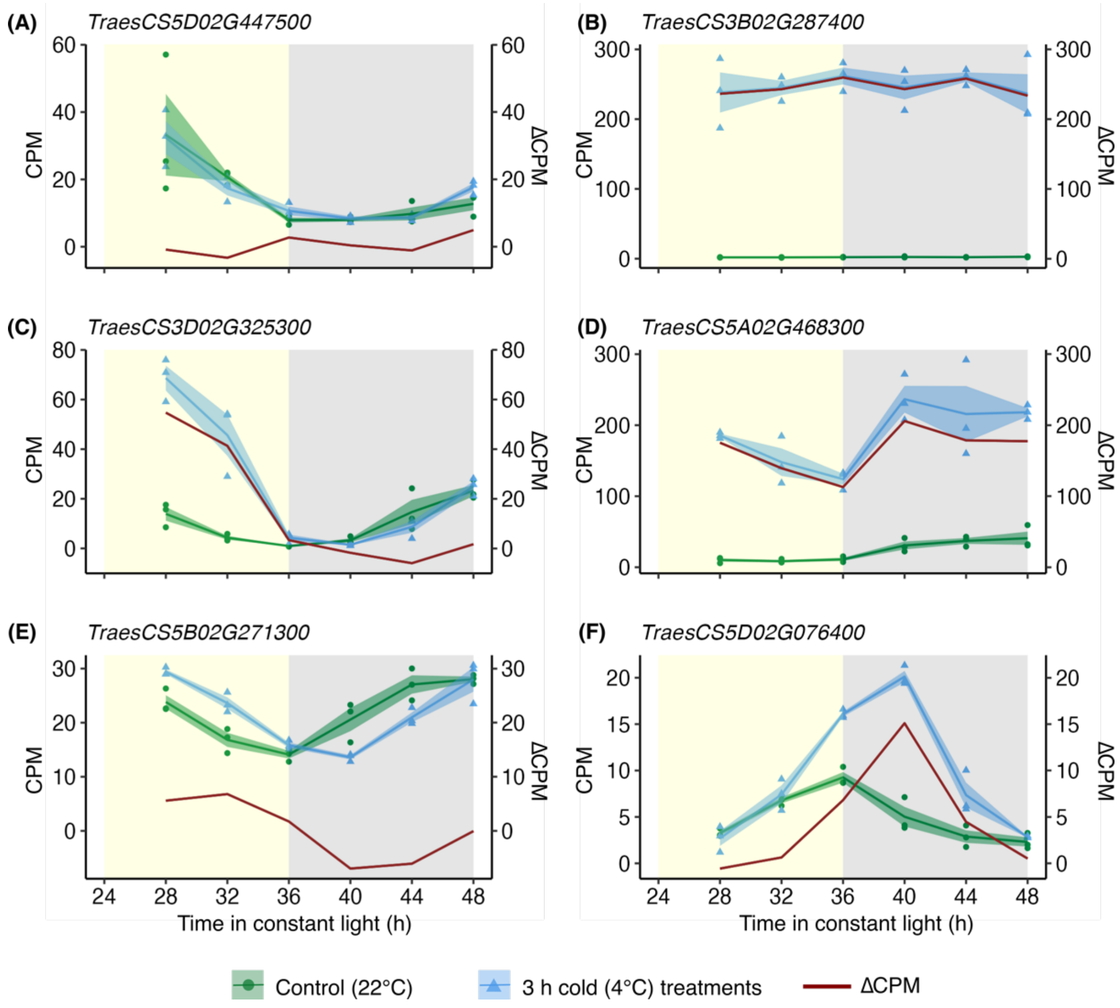


**Figure S8**. **Transcripts exemplifying regulatory features described in Figure 6.** Transcripts **(A)** *TraesCS5D02G447500*, **(B)** *TraesCS3B02G287400*, **(C)** *TraesCS3D02G325300*, **(D)** *TraesCS5A02G468300*, **(E)** *TraesCS5B02G271300*, **(F)** *TraesCS5D02G076400*. Yellow/grey shading = subjective day/night. Solid lines are mean (N = 3 biological replicates). Blue/green shading = ± s.e.m. Red line represents ΔCPM.

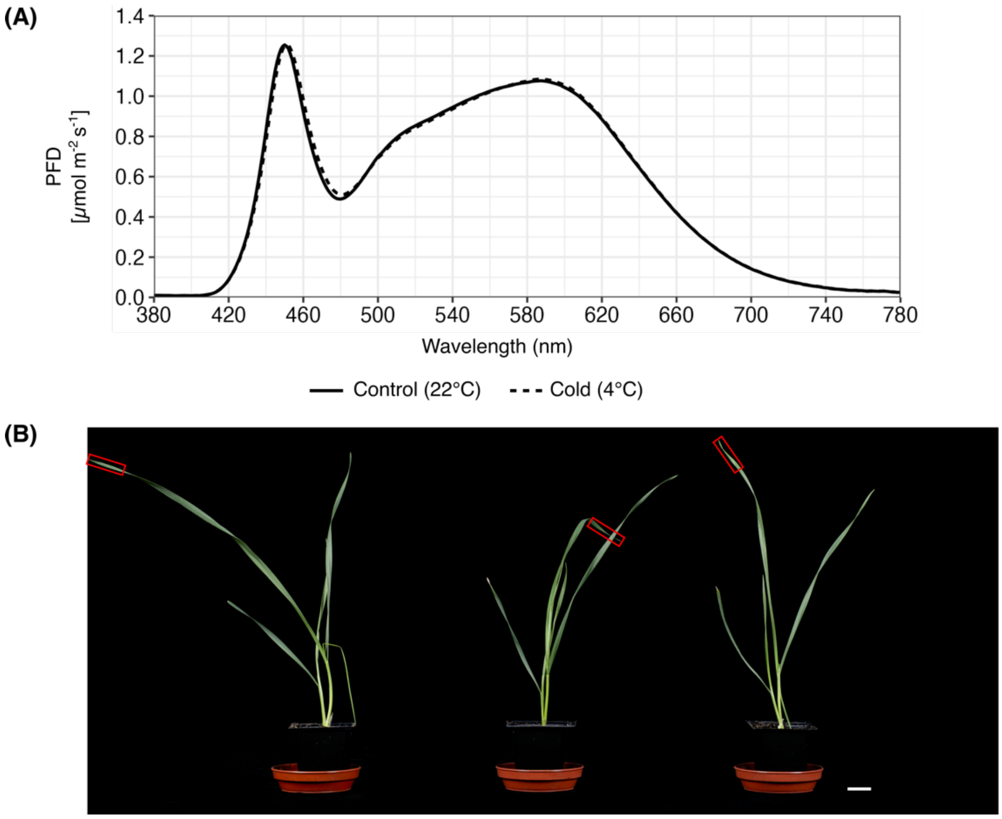


**Figure S9. The light spectra used for cultivation and appearance of seedlings that were sampled**. (**A)** Light spectra used for plant cultivation and experimentation, for control and cold temperature conditions. (**B)** Representative images of bread wheat seedlings after germination on damp filter paper, followed by 12 days growth on compost under 12 h : 12 h light dark cycles (22°C) followed by 24 h of constant light. Seedlings were equivalent to Zadok Stage GS1.2 at time of sampling. Red boxes indicate region of second leaf that was sampled. Scale bar 20 mm.

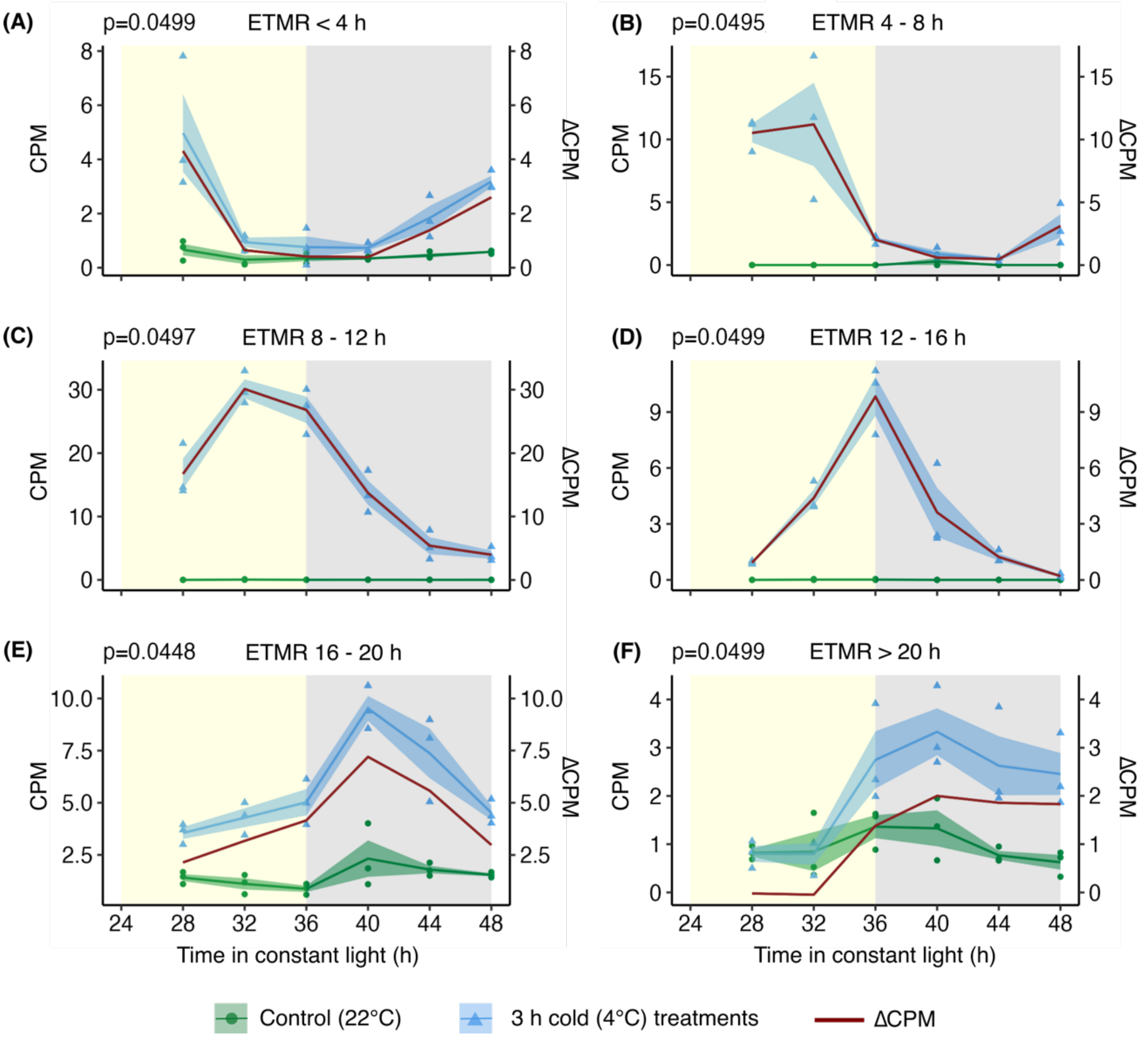


**Figure S10. Transcripts at the upper limit of meta2d statistical threshold for rhythmicity are candidate genes with circadian gating of their response to cold.** Example transcripts that have rhythmic ΔCPM (meta2d $p < 0.05$) and p-value close to the cut-off limit, demonstrating the level of statistical stringency was appropriate for detection of circadian-gated transcripts. Transcripts are from ETMR grouping **(A)** < 4 h, *TraesCS1A02G210400*, **(B)** 4 – 8 h, *TraesCS5D02G343300*, **(C)** 8 – 12 h, *TraesCS5D02G318100*, **(D)** 12 – 16 h, *TraesCS5A02G311100*, **(E)** 16 – 20 h, *TraesCS3B02G342936*, **(F)** > 20 h, *TraesCSU02G072000*. Yellow/grey shading = subjective day/night. Solid lines are mean (N = 3 biological replicates). Blue/green shading = ± s.e.m. Red line represents ΔCPM.
